# A host shift as the origin of tomato bacterial canker caused by *Clavibacter michiganensis*

**DOI:** 10.1101/2023.07.24.550321

**Authors:** Alan G. Yañez-Olvera, Ámbar G. Gómez-Díaz, Nelly Selem-Mojica, Lorena Rodríguez-Orduña, José Pablo Lara-Ávila, Vanina Varni, Florencia Alcoba, Valentina Croce, María Inés Siri, Clavibacter Consortium, Francisco Barona-Gómez

## Abstract

*Clavibacter*, a plant endophytic actinobacterial genus, includes phytopathogens with devasting effects on several crops. *C. michiganensis*, the seed-borne and causal agent of bacterial canker in tomato, is arguably the most notorious species of the genus. Yet, its origin and natural reservoirs remain elusive. Moreover, *C. michiganensis* populations show different genetic pathogenicity profiles with equally unpredictable plant disease outcomes. To tackle these uncertainties, here we analyze genomic data generated during a decade-long survey of *Clavibacter* in wild and commercial tomato cultivars, providing evolutionary insights that informed on the pathogenicity of this phytopathogen. Unexpectedly, our phylogeny situate the last common ancestor of *C. michiganensis* next to *Clavibacter* isolates from grasses rather than to the sole strain we could isolate from wild tomato, which is closer to *C. capsici* associated with pepper. Pathogenicity profiling of selected *C. michiganensis* isolates, together with *C. phaseoli* and *C. californiensis* as sister taxa of the grass clade, and the newly isolated *C. capsici* from wild tomato, was found to be congruent with the proposed phylogenetic relationships. Furthermore, we identified gene enrichment after an evolutionary bottleneck leading to the appearance of *C. michiganesis*, including known pathogenicity factors but also hitherto unnoticed genes with such potential, *i.e.,* nutrient acquisition and specialized metabolite metabolic gene clusters. The holistic perspective provided by our long-term and in-depth analyses hints towards a host shift event as the origin of the causative agent of bacterial canker in tomato, leading to a complex of *C. michiganensis* with pathogenicity factors that remain to be characterized.

## Introduction

Many aspects of human activity are related with the emergence of new pathogens (Sabin et al., 2020). Phenomena like climate change, human invasion of ecosystems and urbanization, can break ecological barriers and create paths that potential pathogens can use to reach new niches (Zhang et al., 2022). Agriculture is no exception: monocultures, as well as high-density cultivation, have been linked to the emergence of new phytopathogens (Stukenbrock & McDonald, 2008), as these practices impose new selective pressures that may favor virulent lineages (McDonald & Stukenbrock, 2016). The introduction of crops to novel environments is another factor that may cause the emergence of pathogens (Weisberg et al., 2021), as this can be transmitted from wild to domesticated hosts, even from unrelated species (McCann, 2020). During this process host shifts take place (Thines, 2019), representing the origin of an emerging disease.

Bacteria of the *Clavibacter* genus, which belong to the phylum Actinomycetota, are mainly known for their phytopathogenic members capable of generating severe diseases in crops of great economic importance (Eichenlaub et al., 2006; Mansfield et al., 2012). The *Clavibacter* phylogeny originally included six subclades referred to as subspecies, which have now been recognized as species (Li et al., 2018) with several new species being proposed (Arizala et al., 2022; Osdaghi et al., 2020). Amongst these, *C. michiganensis* has received a great deal of attention as it is responsible for bacterial canker disease in tomato, *Solanum lycorpersicum* (Sen et al., 2015), with devastating consequences (Poysa, 1993; Mansfield et al., 2012). Even though it was first isolated more than a century ago from tomato crops in Michigan, USA (Smith, 1910), molecular markers are the sole means to tackle the infection, yet reproducibility has been shown to be an issue (Thapa et al., 2020; Yasuhara-Bell et al., 2014). The latter may reflect the fact that genetically diverse *C. michiganensis* populations exist around the globe (Kleitman et al., 2008; Tancos et al., 2015; Valenzuela et al., 2021; Wassermann et al., 2020), and the fact that the origins and natural reservoirs of this phytopathogen remain unknown.

It was not until recently that researchers began to describe genetic features that make this bacterium pathogenic. For instance, Meletzus and co-workers (Meletzus et al., 1993) found evidence that plasmids harbored by *C. michiganensis* contained sequences that encode virulence factors: *celA* on the pCM1 plasmid, which encodes a endo-β-1,4-glucanase (Jahr et al., 2000), and *pat-1* on pCM2, which encodes a serine protease (Dreier et al., 1997). Genome sequencing of the reference NCPPB 382 strain (Stork et al., 2008) allowed the identification of a pathogenicity island (PAI) present in the chromosome. Further genomic studies have shown that *C. michiganensis* has unique features that could explain its pathogenic character in tomato. For instance, Thapa and co-workers (Thapa et al., 2017) showed that certain CAZymes (Carbohydrate-Active Enzymes) families involved in cellulose and hemicellulose degradation are abundant in the *C. michiganensis* genome. Although these previously identified pathogenicity factors have been unambiguously shown to be related to the development of the disease, there is also evidence that some of these factors are absent from strains capable of causing symptoms of the disease (Oh et al., 2022; Tancos et al., 2015) and that virulent strains can infect tomato plants without leading to symptoms (Gitaitis, 1991). Together, these observations suggest that other factors are required for triggering the disease and symptom development (Sharabani et al., 2013, 2014).

Comparative genomics of *C. michiganensis* and other members of its genus has shown that while each species has acquired unique characteristics (Tambong, 2017), there are several features linked to pathogenicity shared between *C. michiganensis* and other *Clavibacter* strains, including cases without background or an antecedent of pathogenicity (Osdaghi et al., 2020). Nevertheless, genomic comparisons have overlooked the relationship between their hosts from which they were isolated and the genomic features that these bacteria may have acquired in their path to adapt to their hosts and develop a pathogenicity lifestyle. To tackle these limitations, here we perform a comprehensive phylogenomics and pangenomic analysis of the *Clavibacter* genus, with an emphasis on *C. michiganensis* strains isolated throughout a decade in geographically distantly and unrelated greenhouses in North (Mexico) and South (Uruguay) America, as well as Europe (the Netherlands). We provide evidence for *C. michiganensis* species sharing a common ancestor with isolates from different grasses, and that the species has undergone an evolutionary bottleneck. Our analysis revealed genetic markers, with functional implications *in planta*, that evidence a host shift, from grasses to tomato. We provide an example on how to track the origin of emerging plant pathogens and pave the way to further functionally characterize the pathogenicity of *C. michiganensis* at the population level in line with its dual symptomatic and asymptomatic behavior.

## Methods

### Bacterial strain isolation and identification

Tomato (*Solanum lycopersicum*) plants with symptoms but also asymptomatic were collected during 2010 to 2020 from several geographically distant sites in Mexico, mainly from high-tech greenhouses (Aguascalientes, Colima, Guanajuato, Michoacán, Nuevo León, Querétaro, San Luis Potosí and Zacatecas), and from two wild populations (Jalisco and Guanajuato). Samples consisted of stem, leaves, and fruit when available. Tissue was cut into slices of approximately 0.5 cm^2^ of surface area and placed directly into petri dishes with CMM1 semi-selective media (for 1 liter: 10.0 g saccharose 1.2 g Tris base, 250 mg MgSO_4_·7H_2_O, 5.0 g LiCl, 2.0 g yeast extract, 1.0 g NH_4_Cl, 4.0 g casamino acids and 15.0 g agar; supplemented with 28 mg/l nalidixic acid, 10 mg/l polymyxin B sulfate and 200 mg/l cycloheximide). CMM1 Petri dishes were incubated at 28°C and growth monitored for 24 to 72 hrs. Grown colonies were selected based on *C. michiganensis* reported morphology (EPPO, 2016) and isolated and cultured on individual petri dishes with CMM1 media. LB liquid cultures were prepared for each isolate for DNA extraction, carried out with alkaline lysis, phenol-chloroform extraction, and ethanol precipitation. *C. michiganensis* isolates were identified by PCR using the *clvF* marker gene (Yasuhara-Bell et al., 2014). *clvF* PCR negative isolates were identified by Sanger sequencing of the PCR amplified 16S rRNA gene. A total of 550 plant specimens, leading to a strain collection of around 151 *Clavibacter* positive isolates, were obtained (148 *C. michiganensis* confirmed strains).

### Genome sequencing and database

DNA from the 151 *Clavibacter* positive strains was extracted, as previously stated, for whole genome sequencing using Illumina paired-end MiSeq or NextSeq platforms. Read quality was assessed with FastQC. Poor quality sequences were trimmed with Trimmomatic (Bolger et al., 2014), with values adjusted for each read set. *De novo* assembly was performed with the smart and auto assembly strategies in PATRIC (Davis et al., 2020). Seven representative genomes of the *C. michiganensis* strains isolated in Mexico from commercial cultivars were chosen based on independent phylogenomic and pangenome analysis made at species level followed by a selection of those strains that best represented the genomic diversity according to the gene families present in their respective clades (data unpublished).These selected genomes and 58 publicly available genomes were used to construct an *ad hoc* genomic database (DB) which was complemented with *C. michiganensis* genomes obtained from strains isolated from tomato cultivars with symptoms from other parts of the world: 3 from Uruguay and 1 from the Netherlands. From the total of 69 *Clavibacter* genomes that comprised the complete *Clavibacter* genus DB, 32 corresponded to *C. michiganensis* sequences. All genome assemblies comply with quality metric thresholds: ≥25000 for the N50 and ≤36 for L50. The RAST tool kit was used for gene calling and function prediction (Brettin et al., 2015), complemented with the Pfam database (Mistry et al., 2021).

### Clavibacter phylogeny and pangenome analysis

Single copy core genes in the previous DB were identified using Anvi’o (v7.0) pangenome analysis tool (Eren et al., 2021). Identical genes were filtered out based on the functional homogeneity index (<0.999). A total of 1231 genes were selected. Aligned and concatenated amino acid sequences were obtained with *anvi-get-sequences-for-gene-clusters*. The concatenated sequences were used to infer phylogenetic relationships by generating phylogenetic trees with IQ-TREE 2 (Minh et al., 2020). Branch support was assessed using the ultrafast bootstrap approximation from UFBoot (Hoang et al., 2018) with 1000 replicates. Best-fit substitution model was determined using ModelFinder (Kalyaanamoorthy et al., 2017) restricting the testing procedure to the WAG (Whelan & Goldman, 2001) and LG (Le & Gascuel, 2008) models. Branches of the *Clavibacter* genus phylogeny were ordered using a family-level reference tree, which included 15 *Clavibacter* strains and *Rathayibacter toxicus FH232* as a root (**Supp. Fig. 1**). A pairwise Average Nucleotide Identity analysis of all the genomes was performed with *anvi-compute-genome-similarity* using PyANI (Pritchard et al., 2015) with BLAST and default values. The DB was analyzed using the program *anvi-pangenome* from Anvi’o, using default parameters unless stated otherwise. Briefly, the program calculated the similarity between amino acid gene sequences with BLASTp (Altschul et al., 1997), then resolved gene families with the MCL algorithm (Van Dongen & Abreu-Goodger, 2012) using an inflation value of 10. Eren et al. (2021) refer to *gene clusters*, which we call *gene families* in this manuscript, to avoid confusion when referring to Biosynthetic Gene Clusters (BGCs).

### Identification of genomic evolutionary signals

Gene family enrichment analysis was performed at the genus-level with the cognate genome DB using anvi’o’s *anvi-compute-functional-enrichment* program and the *–include-gc-identity-as-function* option. Briefly, this program groups the genomes in the dataset according to a categorical variable, in our case membership to the so-called “*C. michiganensis* broad clade”, and then determines which gene families are enriched in the genomes within our group of interest and absent, or nearly absent, in the rest of the genomes. The statistical approach to determine the enrichment was described elsewhere (Shaiber et al., 2020). Parallelly, we looked for phylogenetic signals in the occurrence of the identified gene families with the D measure (Fritz & Purvis, 2010) using caper (v. 1.0.1) R package (Orme et al., 2013). D was estimated for each gene family using the *Clavibacter* genus phylogeny and the presence and absence of the gene families in each genome as the binary trait to evaluate. Gene families were considered enriched when their enrichment score was ⋝50, their q-adjusted value from the enrichment analysis was < 1e-10 and the value of D was < 0. For the identification of evolutionary and functionally informative loci, genes from the reference strain NCPPB 382 occurring in the enriched gene families were extracted using Anvi’o pangenome analysis tool. Since this is a closed genome, their annotation order corresponds to their position in the chromosome, hence genes were ordered accordingly and checked for contiguity. Only genes with at least three enriched neighboring genes were considered for further inspection. Genes were still considered neighbors even when there was a gap of maximum two non-enriched genes between them. Genes within the pathogenicity island (PAI) were identified according to their location relative to the genes that limit this region as reported by Gartemann et al. (2008).

### Evolutionary genome mining of natural products

Given the results obtained during the identification of evolutionary genomic traits, we focused on the prediction and analysis of BGCs encoding for Ribosomally synthesized and Post-translationally modified Peptides (RiPPs). RiPP type BGCs were searched in the genome DB using antiSMASH v6.0 (Blin et al., 2021), including RRE-Finder (Kloosterman et al., 2020) to improve RiPP BGC identification. Identified RiPP BGCs were grouped into similarity networks using BiG-SCAPE (Navarro-Muñoz et al., 2020) employing a cut off value of 5.0 and compared against version 3.0 of MiBIG database (Terlouw et al., 2022). Selected BGCs were analyzed further with CORASON to solve their evolutionary relationships at the whole BGC-level (Navarro-Muñoz et al., 2020). Predicted structures were based on previously reported literature (Holtsmark et al., 2006; Wiebach et al., 2018) and are provided only as a proxy to chemical classes involved, but remain to be experimentally confirmed. CORASON (Navarro-Muñoz et al., 2020) was used to explore the genomic neighborhood around the RiPP BGCs in *C. michiganensis* NCPPB 382 to identify homologue genes in other *Clavibacter* species. Using antiSMASH’s annotation of the RiPP BGCs as a reference, the amino acid sequences of non-biosynthetic genes were used as query for CORASON increasing the *cluster_radio* value to 30. After identifying homologous genes in the BGCs’ genomic neighborhood, Easyfig (Sullivan et al., 2011) and BLASTn (Zhang et al., 2004) were used to compare the neighboordhood’s sequences at the nucleotide level. CORASON’s output was used as a reference to color homologous genes in Easyfig’s output figure.

### Phenotypic characterization in planta of C. michiganensis genetic diversity

Disease development assays were performed at a greenhouse in July 2020, June 2021 and August 2021. *Clavibacter-*free tomato plantlets were provided by a commercial supplier (Plantanova, Mexico) with three to four true leaves, which were inoculated with selected *C. michiganensis* strains from our collection. Three non *C. michiganesis* strains, namely, *C. californiensis* CFBP 8216, *C. phaseoli* CFBP 8217 (both acquired from the corresponding type strain collection) and *C. capsici* RA1B (isolated by us from wild tomato) were also included. Bacterial inoculation was made by scraping the surface (0.5 cm^2^) of the stem with a needle below the first two true leaves and then puncturing slightly at the center of the scrapped area. Inoculum consisted of 5 µl of bacterial culture with a concentration of 1.5×10^9^ CFU. Mock-inoculated control plants were generated using sterile water. Disease development was monitored in a weekly fashion for 6 weeks after inoculation. Disease progression classification and disease index calculation were performed as in Valenzuela et al. (2021) for each week to obtain a disease progress curve. The so-called Area Under Disease Progress Curve (AUDPC) was calculated using the *audpc* function from the agricolae package in R (Mendiburu, 2022).

## Results

### C. michiganensis prevails in commercial tomato cultivars but not in wild relatives

To our knowledge, tomato bacterial canker has only been reported in commercial tomato cultivars, which led us to search for *C. michiganensis* existing in tomato wild relatives. If so, these wild plants could act as a natural reservoir for pathogens jumping to related host plants. Moreover, if they existed, these bacteria could provide us with valuable insights on the evolution of *C. michiganensis* to adapt and thrive in tomato cultivars. Hence, we attempted to isolate *Clavibacter* strains from wild tomato populations. We sampled 39 plants from two different populations and isolated 222 bacterial strains on semi selective CMM1 medium, which has been reliably used for isolation of *C. michiganensis* (Alvarez et al., 2005). Based on their 16S rRNA gene, most of the strains isolated correspond to the *Micrococcales* bacterial order (**Supp. Table S1**). Unexpectedly, only one isolate, termed RA1B, belonged to the *Clavibacter* genus, which came from a wild tomato plant with no noticeable disease symptoms. The 16S rRNA indicated that RA1B belongs to the *Clavibacter* genus, while ANI (Average Nucleotide Identity) analysis performed after genome sequencing showed that RA1B resembles *C. capsici*: 90.29% percentage identity when compared with *C. michiganensis* NCPPB 382 *vs.* 98.93% when compared with *C. capsici* PF008 (**Supp. Fig. 2, Supp. Table S2**).

### *Clavibacter* has undergone multiple hosts shifts unrelated to wild tomatoes

To determine the phylogenetic relationships among our strains and other previously reported *Clavibacter* strains, we assembled a genus DB including diverse *Clavibacter* genomes (**Supp. Table S3**). Then, we reconstructed a phylogenomic tree using 1231 single copy genes found to be conserved in the 69 genomes. Even though *C. michiganensis* is commonly associated with tomato, the fact that we managed to isolate a *C. capsici* strain from a wild tomato variety, prompted us to analyze the relationship between the bacterial phylogeny and the hosts from which they were isolated (**Fig. 1**). Clades and subclades were defined based on the tree topology and ANI values (**Supp. Fig. 2**, **Supp. Table S2**), where monophyletic strains with >97% and >93% ANI similarity were considered part of the same subclade or clade, respectively. Thus, *Clavibacter* strains grouped into ten different subclades, plus eleven single-strain lineages (SSLs) that remain to be populated with further isolates and their genome sequences.

The phylogeny revealed that *Clavibacter* strains come from a more diverse and broader host diversity than expected. Moreover, the host-strain relationship seems to be independent of the evolutionary history of the strains, as a lack of relationship between the species, their pathogenicity, or the position of each species in the tree, was found. For instance, the *Clavibacter* species that are pathogenic in plants of the same family do not cluster together but are situated in distantly related clades: *C. nebraskensis*, pathogenic in maize (a *Poaceae* plant) and *C. insidiosus*, pathogenic in alfalfa (a *Fabaceae*) are clustered into sister clades. In contrast, *C. capsici, C. sepedonicus* and *C. michiganensis*, which infect *Solanaceae* plants, are placed at early and late divergent subclades. Moreover, *C. michiganensis* and *C. sepedonicus* both diverged after a clade of *Poaceae* isolates we termed the Pre-Solanaceae Jump (PSJ) clade, since their ancestor diverged from the lineage that precedes both *Solanaceae*-related species. SSLs with no reported pathogenic activity (1, 4, 5, 6, 7, 10) form monophyletic clades with the pathogenic strains. SSLs 4 and 5 cluster with *C. insidiosus* and *C. nebraskensis* subclades, SSLs 6 and 7 cluster with *C. phaseoli* (formerly known as *C. chilensis*, Arizala et al., 2022) while SSL10 cluster with SSL9 and the *C. tessellarius* sub-clade. The case of the so-called “Broad *C. michiganensis* clade” (from now onwards BCm clade) is particularly interesting since the *C. michiganensis* pathogenic subclade clusters with three non-pathogenic groups: SSL1 and the Pre-Phytonotic event 1 and 2 subclades (PP1 and PP2, **Fig. 1**), which were named as such since they descend from the lineage that diverged from the one that originated *C. michiganensis*.

Except from the *C. nebraskensis* subclade, all the subclades consist of strains obtained from at least two different host species. While there are cases where the host species are closely related, even belonging to the same plant genus (e.g., *C. insidiosus* strains), there are several others where strains grouping together come from distantly related hosts, such as in the *C. tessellarius, C. phaseoli* and *C. michiganensis* subclades. Notably, several *Clavibacter* strains isolated from *Poaceae* and *Solanaceae* plants (particularly from tomato) are distributed throughout several clades. In some cases, strains isolated from both plant families clustered within the same clade, such as in the so-called “Early Divergent clade” (from now on ED clade), and in the *C. tessellarius, C. phaseoli* and *C. michiganensis* subclades (**Fig. 1**). This pattern is particularly noticeable in the subclade of *C. michiganensis* species, which includes several cases of host shifts. This large clade, whose richness reflects the fact that we and others have sampled it thoroughly, includes several strains obtained from other plants other than tomato, with many *Poaceae* host plants examples but not the RA1B isolate from wild tomato.

**Figure 1.**
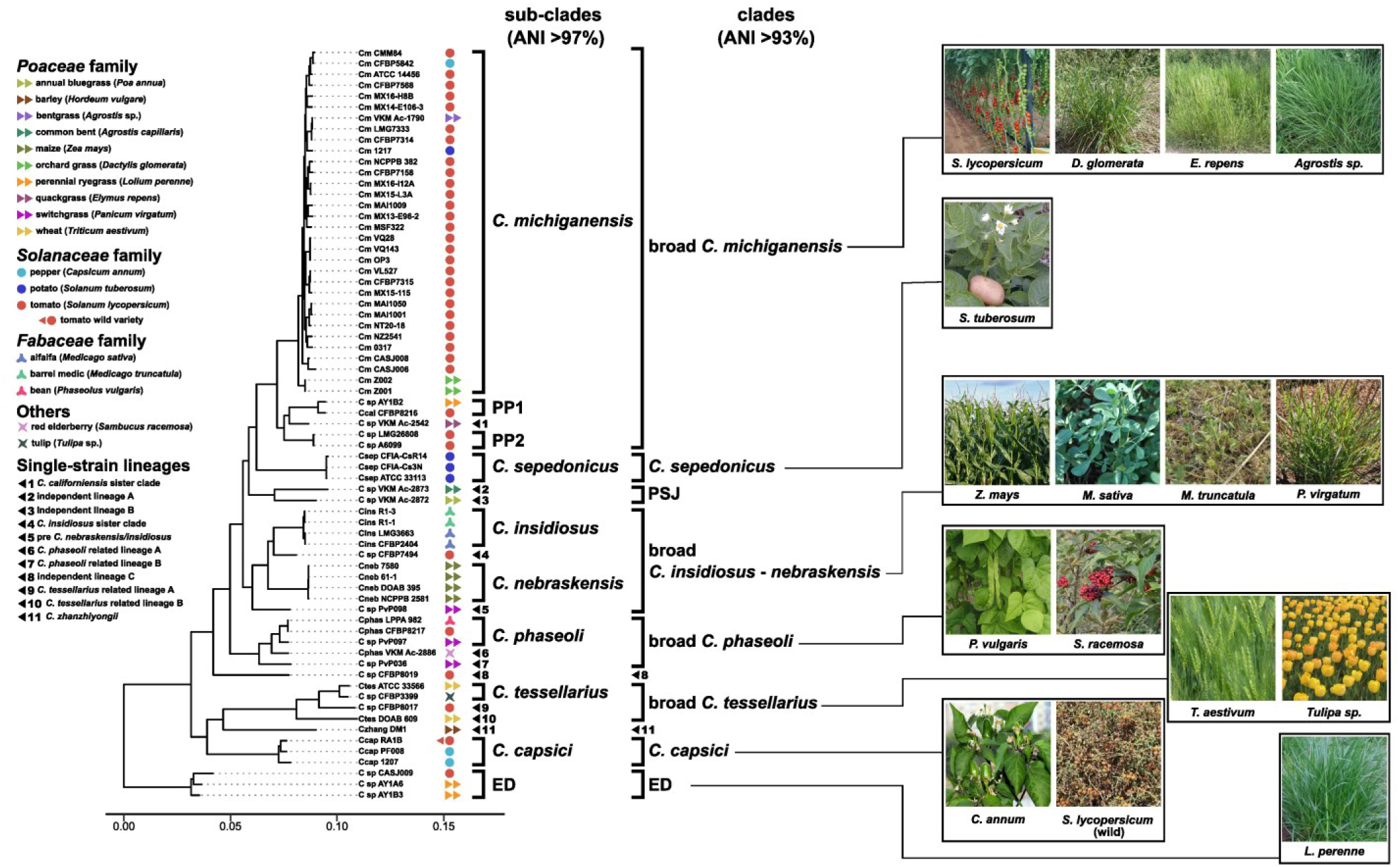
Phylogenomic tree of the *Clavibacter* genus and hosts of origin. Left: phylogenomic tree with clades and lineages indicated with brackets and numbers. Each host species is indicated by the colored geometrical shapes next to the strains’ names. Right: representative pictures of plant hosts from which strains were isolated. Tree was inferred from 1231 core proteins. Only posterior probability support values <0.9 are shown. *Cm, C. michiganensis; C sp., Clavibacter* sp.; *Ccal, C. californiensis*; *Csep*, *C. sepedonicus*; *Cneb*, *C. nebraskensis*; *Cins, C. insidiosus*; *Cphas*, *C. phaseoli*; *Czhang, C. zhanzhiyongii*; *Ccap, C. capsici*; *Ctes, C. tessellarius*. ED = Early Divergent. PP1 = Pre-phytonotic event-1. PP2 = Pre-phytonotic event-2. PSJ = Pre-*Solanaceae* Jump.

### Pathogenicity profiles of selected strains supports the phylogeny and host shifts

To test the phenotypic implications of our phylogeny, we then assessed the pathogenicity of *C. capsici* RA1B in tomato plants and compare it to the two most highly pathogenic strains of our *C. michiganensis* collection, strains MX15-115 and MX16-I12A. Moreover, with these experiments, we also aimed at contrasting the pathogenicity profiles of these known pathogens against the non michiganensis *Clavibacter* strains isolated from tomato: *C. californiensis* CFBP 8216 and *C. phaseoli* CFBP 8217, previously reported to be asymptomatic (Yasuhara-Bell & Alvarez, 2015). These two strains, which provided a proxy to the grass strains (not available to us), were referred to as the not-michiganensis tomato *Clavibacter* (NMTC) and belong to the PP1 and *C. phaseoli* subclades (**Fig. 1**). These *Clavibacter* strains were treated equally and used to infect tomato plants to measure the development of disease symptoms, which were monitored for a period of six weeks (**Fig. 2**). Throughout the period of evaluation, RA1B did not show any sign of being pathogenic in tomato plants. Surprisingly, NMTC isolates showed the development, even if very mild but yet quantifiable, of clear symptoms compared to the *C. michiganensis* strains.

**Figure 2.**
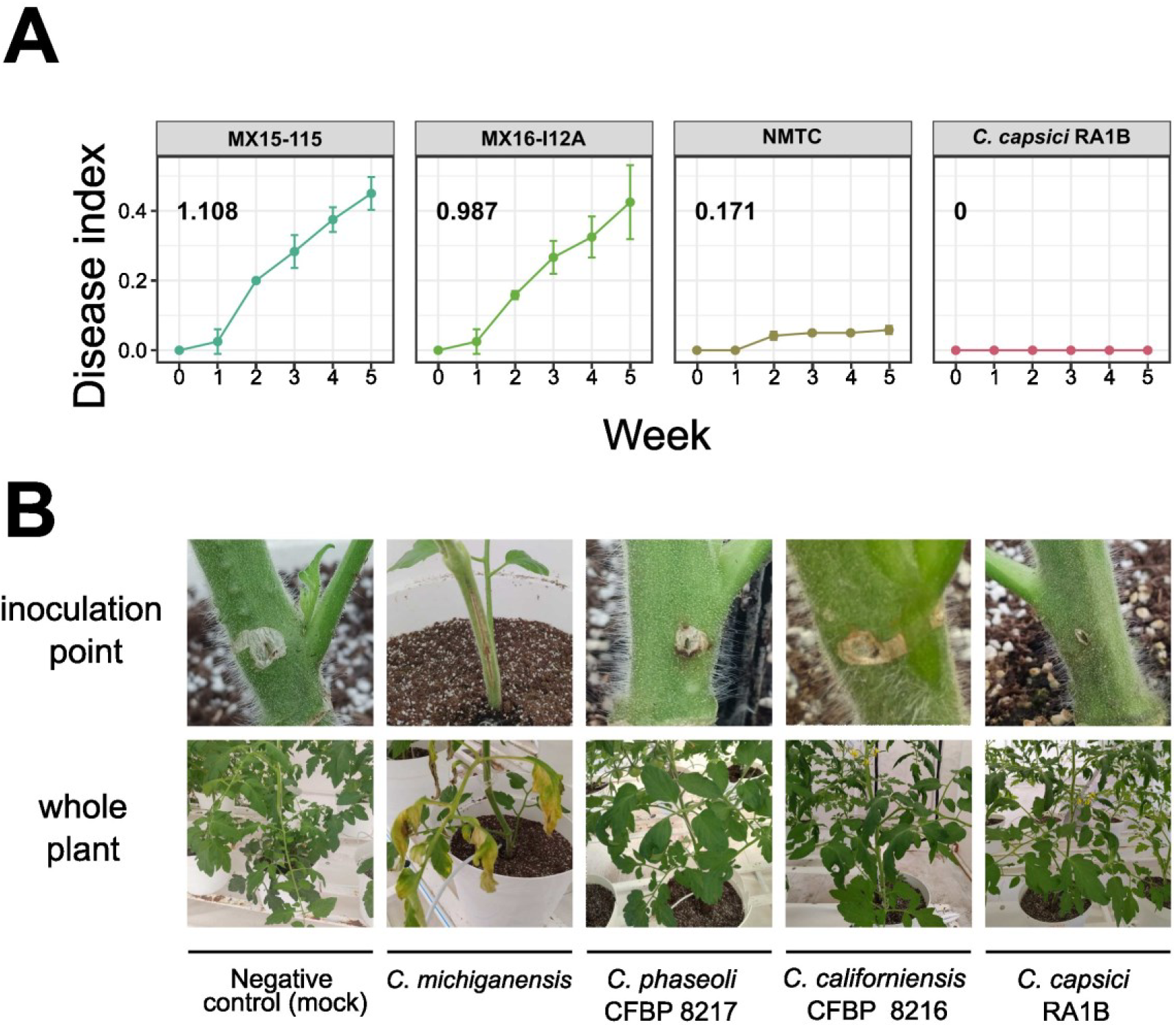
Pathogenicity profiling of selected *Clavibacter* strains. **A.** Pathogenicity profiling was quantitively done for selected strains using a disease index calculated for each treatment per week and used to calculate the area under disease progress curve (numbers in bold at top right of each subplot). **B.** Qualitative comparison of symptoms generated by selected *Clavibacter* strains, including *C. michiganensis* (strong symptoms), NMTC strains *C. phaseoli* and *C. californiensis* (mild symptoms only in the site of inoculation) and *C. capsici* RA1B (asymptomatic).

### Genetic linkages between grass and commercial tomato isolates indicative of a host shift event

When colonizing a new host, pre-adapted pathogens undergo changes that allow them to better thrive in their new environment. To identify these features at the genome level, we compared the gene content of key strains throughout the *Clavibacter* genus, and searched for genes that were associated with *C. michiganensis*. To achieve this, we performed a pangenomic analysis using the *Clavibacter* genus DB previously assembled. This analysis identified 11447 different gene families, of which 1766 were shared among all the strains in the dataset and 4183 were unique to one strain (singletons). Since our aim was to identify features common only to the *C. michiganensis* strains, conserved gene families and singletons were discarded, while the analysis continued with the 5498 remaining gene families. We then identified gene families acquired at the split of the *C. michiganensis* subclade and its sister clades within the BCm clade, as this would be the time of the occurrence of the proposed host shift. To test this hypothesis in more detailed, we employed two complementary approaches, as described next.

First, a gene enrichment analysis using Anvi’o was adopted to identify gene families enriched in the BCm clade and absent (or nearly absent) outside of it. This allowed us to filter out gene families shared by the strains belonging to the BCm clade from other clades of the *Clavibacter* genus. As most of the BCm clade consists of *C. michiganensis* strains, we expected that the gene families with the highest enrichment scores would be those with high degree of conservation in this species. Second, we identified gene families whose presence and absence pattern in the genus phylogeny had a phylogenetic signal. This allowed us discard gene families whose presence, although enriched in the BCm clade, were randomly distributed across it because of natural genomic variation. For this, we use the presence and absence of the gene families in the genomes as a binary trait whose phylogenetic signal could be tested under Fritz and Purvis’ D. Conjunctly, these two approaches allowed us to identify 103 gene families (**Supp. Table 4**) that were conserved in the BCm clade (**Fig. 3**). We further classified the conserved gene families between those that were present outside the *C. michiganensis* clade from those that were found exclusively in these species.

Operonic gene organization in bacteria is suggestive of functional association between genes that cluster together on the same genomic region. For this reason, we analyzed if the genes identified as conserved gene families co-locate in the genome of *C. michiganensis*. Using strain NCPPB 382 as a reference, we defined these genes’ location, leading to the identification of nine regions or loci (labeled from 1 to 8, and PAI) consisting of neighboring genes (**Fig. 3**, left). The first and largest region identified corresponds to the pathogenicity island (PAI) identified by Gartemann et al., (2008). Indeed, a total of 29 gene families containing genes associated with the PAI were identified by our analysis. Only three of the identified gene families belonging to the PAI had homologs outside the *C. michiganensis* clade. One of them correspond to a family of hypothetical proteins, while the other two correspond to gene families related to the AbiEii toxin/antitoxin complex, which helps bacteria to deal with phages. Interestingly, genes of these last two gene families can be found, besides the PAI, in the pCM2 plasmid (**Supp. Table 4**). Likewise, loci 5 was found to be the michiganin biosynthetic gene cluster (BGC), known to encode for the synthesis of a ribosomally synthesized and post-translationally modified peptide (RiPP) previously isolated (Holtsmark et al., 2006) and used for PCR diagnostic purposes (Yasuhara-Bell et al., 2014). These loci belong to the gene families with some occurrence outside the *C. michiganensis* clade, albeit at a low frequency. Besides the PAI and michiganin BGC, whose identification confirm the validity of the approach adopted, we identified seven more loci indicative of the evolutionary dynamics leading to *C. michiganensis*.

Three loci belonged to genes exclusively found in *C. michiganensis*, as the PAI (i.e., 3, 4 and 8), and four loci included gene families present outside this clade, like the michiganin BGC (i.e., 1, 2, 6 and 7). Functional annotation of the genes conformed these novel *C. michiganensis* loci and predicted their putative roles during evolution of this pathogen. These genes were found to classify in three different types, (i) transport systems, (ii) RiPP BGCs, and (iii) undetermined (**Fig. 3**, right). Four out of the seven new loci identified (1, 2, 7 and 8) have genes suggestive of transport systems. Genes within loci 2 and 7 were annotated as glycosyl hydrolases, which suggests the transporters encoded in these loci participate in carbohydrate assimilation. Locus 1 was annotated as a Major Facilitator type transporter, but the substrates upon which this system may operate, could not be predicted. For annotation of the hypothetical or genes of unknown function included in locus 8, many of which were found to be conserved in other actinobacteria, we used ProteInfer. These analyses hinted towards nitrogen metabolism, including metabolites such as serine, as possible functions encoded by these genes. Also, an unprecedented RiPP BGC, *i.e.,* locus 4, could be annotated with antiSMASH. We termed the putative product of this BGC ‘michivionin’, as it shares the same chemical class (RiPP class III) with microvionin from *Microbacterium arborescens* 5913 (Wiebach et al., 2018), which is further analyzed in detail at the sequence level in the final section

**Figure 3.**
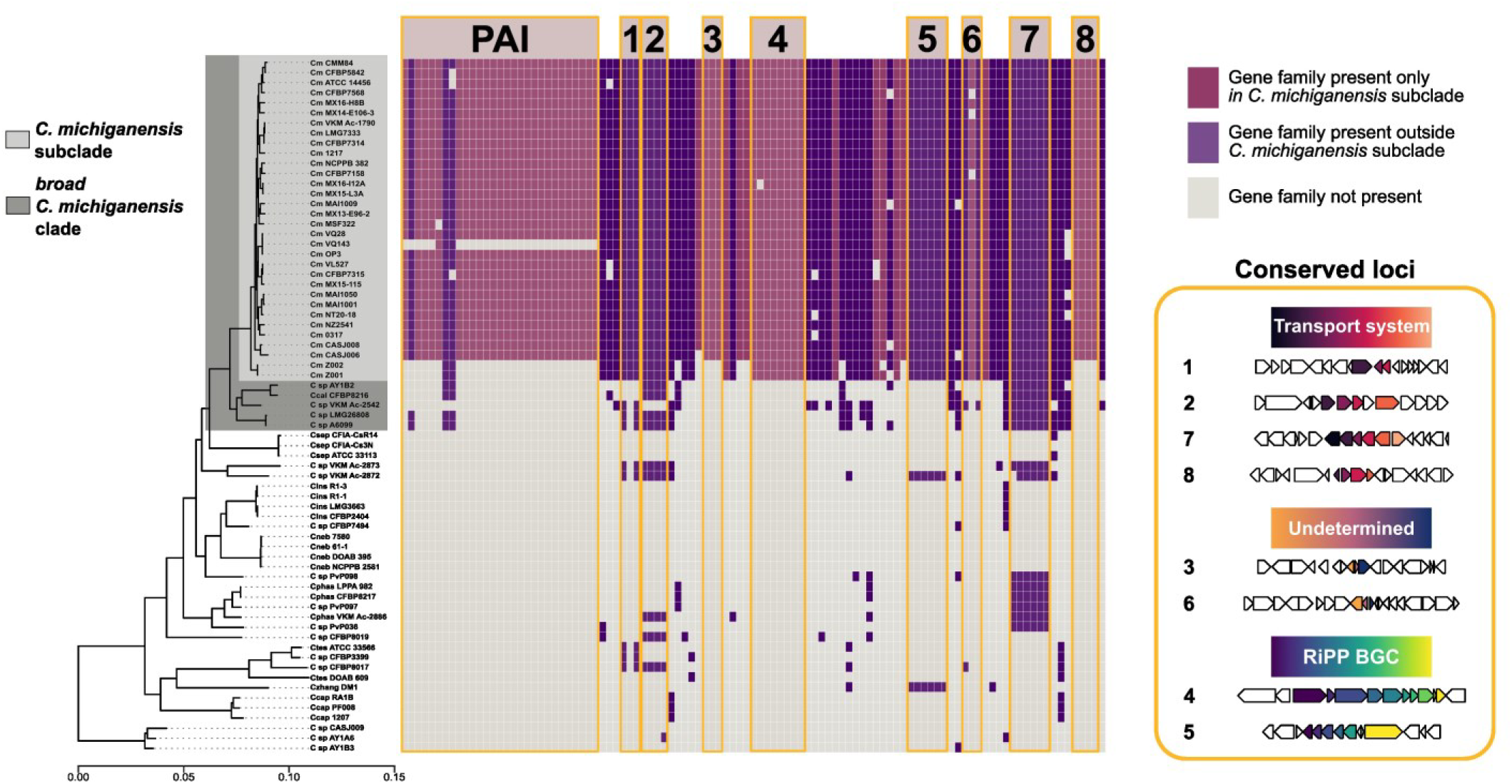
Conserved loci in *C. michiganensis* selected during the proposed host shift. **Left**, Heatmap of characteristic gene families of *C. michiganensis*. Columns of the heatmap represent different gene families in each of the analyzed strains throughout the phylogeny. The highlighted clade (dark gray) shows the BCm clade. Purple columns represent gene families found outside the *C. michiganensis* subclade, while magenta columns indicate gene families present exclusively in *C. michiganensis* subclade. Columns are ordered according to the occurrence of genes in the genome of *C. michiganensis* NCPPB 382. Gene families corresponding to the pathogenicity island are indicated as PAI. **Right.** Identified loci consisting of more than three contiguous, or semi contiguous genes, as described in Methods. Predicted functions of each gene in these loci is displayed below for each region. Associated function for each loci is indicated by a distinctive color gradient: purple è yellow, RiPPs; deep purpule è orange, transport systems; orange è blue, undetermined.

### Genome mining of RiPPs implicated with *Clavibacter* adaptation to tomato

Since the results of the pangenomic analysis suggested that RiPP BGCs may be a characteristic feature of *C. michiganensis*, we decided to explore the biosynthetic potential of the entire genus for these ecologically relevant metabolites (Li & Rebuffat, 2020). For this, we annotated the RiPP BGCs in the genus DB and group them into Gene Cluster Families (GCF) using BiG-SCAPE (**Fig. 4A**, top). We then resolved these GCFs throughout the genus phylogeny (**Fig. 4A**, bottom). Overall, we found twelve networks each one corresponding to a different GCF distributed across the whole genus. Only network I, corresponding to a Linear azol(in)e-containing peptide (LAP) RiPP BGC, was fully conserved throughout the genus. Along with this network, II, IV and IX were also found in two or more clades, *e.g.* network II in the *C. michiganensis* subclade, the SSL3 and in the *C. zhangzhiyongii* strain; network IV in the *C. sepedonicus* and *C. capsici* clades; and network IX in SSL1 and PCC2. Networks III, V and VI were found to be exclusive for one *Clavibacter* species each: *C. michiganensis*, *C. capsici* and *C. sepedonicus*, respectively, although the subclades of the latter two species are poorly populated, which is the case for the rest of the networks with only one strain. In contrast, the presence of networks II and III (**Fig. 4B**, top), which correspond to locus 5 (michiganin BGC) and locus 4 (michivionin BGC) from the previous pangenomic analysis, are what distinguishes *C. michiganensis* from the other *Clavibacter* species.

The previous analysis revealed that the michiganin BGC, once considered a distinctive feature of *C. michiganensis*, is shared with other *Clavibacter* species. Hence, we decided to inspect the genomic neighborhood of this BGC and compare it with other *Clavibacter* species to gain insights about the evolutionary history of this and other BGCs. Even though the michivionin BGC is exclusive to *C. michiganensis*, according to our results, we inspected and compared its genomic neighborhood as well. We selected thirteen *Clavibacter* strains representing main clades of the genus as resolved by the phylogenetic tree and performed a synteny analysis of the surrounding of both BGCs using CORASON (**Fig. 4B**, bottom).

Based on the identified homologous genes in each loci, we then used Easyfig and BLASTn to compare the nucleotide sequences of the different loci. BLASTn alignment for regions over 1000bp showed high degree of similarity for both neighborhoods: 89.68 ± 5.62% identity (e-value 3.18e-70 ± 6.13 e-69) for michiganin’s BGC neighborhood and 90.60 ± 2.92% (e-value 0) for michivionin’s BGC vicinity. Out of the two loci, the michivionin neighborhood is highly conserved in every *Clavibacter* strain. The sole difference was the presence of the michivionin BGC itself. In contrast, genomic neighborhood around the michiganin BGC is more variable, yet orthologous genes could still be identified throughout all strains. Interestingly, a michiganin-like BGC exists in *Microbacterium arborescens* 5913, the microvionin producer, coincidently in the vicinity of its cognate BGC. Comparison was then expanded to include this *Microbacterium* species (**Fig. 4B**), showing sequence and gene organization similarities only for the michiganin-related BGCs but not for the analogous microvionin and michivionin BGCs. Despite these, our analyses suggest a possible correlation between RiPPs BGCs, horizontally transferred amongst related plant *Microccocales* genera, and evolution of *C. michiganensis* as a tomato pathogen from grass endophytes.

**Figure 4.**
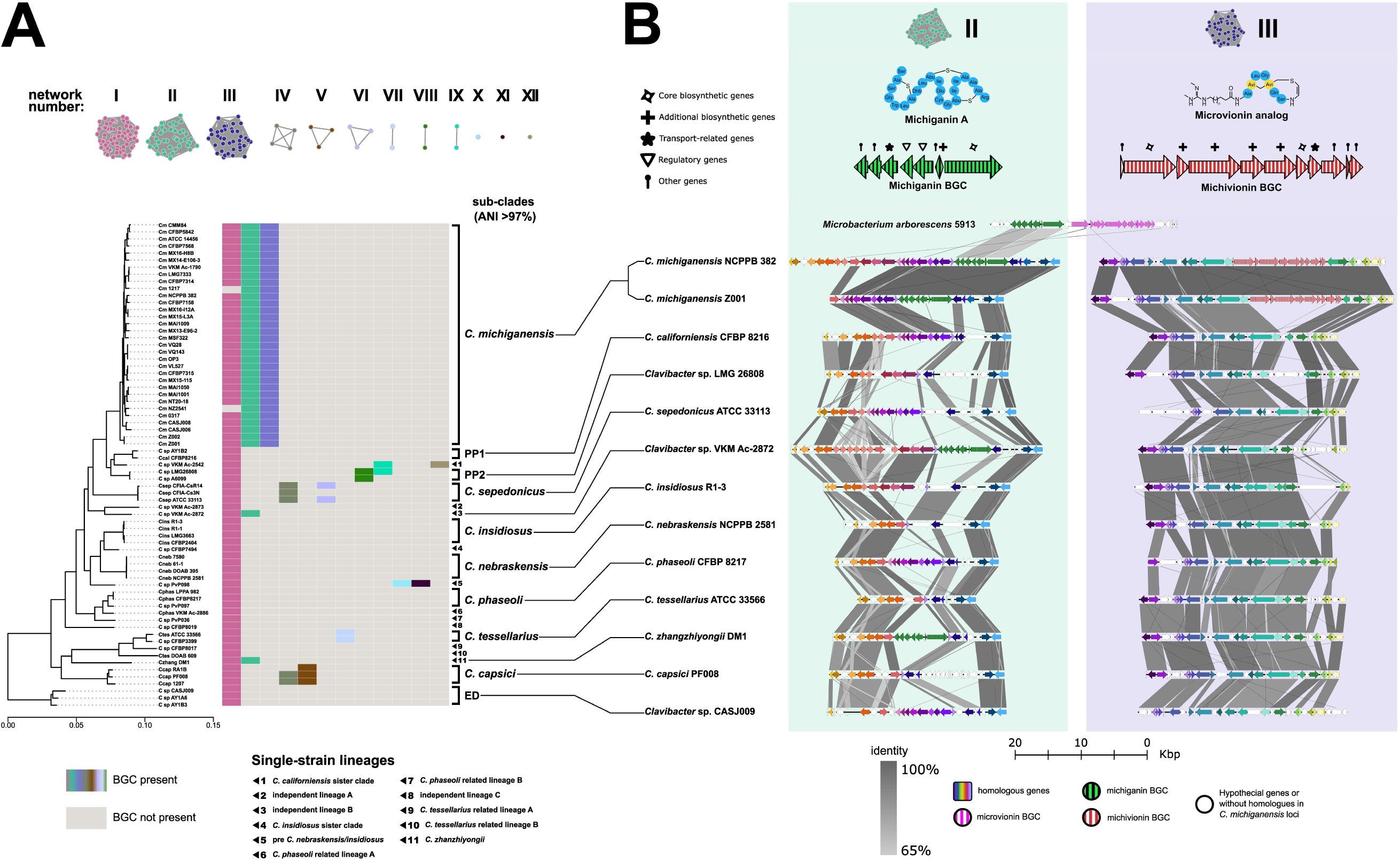
Predicted RiPP BGCs in *Clavibacter*. **A.** Top: RiPP BGC networks representing a different set of GCFs. Bottom: Presence of each BGC network in the *Clavibacter* genus, with each column displaying the presence or absence of each BGC as they appear in the top part of the panel. ED = Early Divergent. PP1 = Pre-Phytonotic event-1. PP2 = Pre-Phytoonotic event-2. **B.** Top: RiPP BGCs specific to the *C. michiganensis* subclade. The BGC from *C. michiganensis* NCPPB 382 is shown. Representative structures of the type of RiPP expected are shown in the right-hand side. Bottom: Genomic neighborhood of michiganin and microvionin BGCs highlighting horizontal gene transfers. The genomic neighborhood of michiganin and michivionin BGCs of *C. michiganensis* NCPPB 382 compared to the same loci in thirteen selected *Clavibacter* strains. Same colors are used for homologous genes believed to be orthologous based on sequence similarity and shared synteny. The microvionin BGC from *Microbacterium arborescens* 5913, which includes a michiganin-like BGC in its vicinity, is displayed on the top of the other loci.

## Discussion

The *Clavibacter* genus has been known for a long time, mainly because of their pathogenic members. With some exceptions, it seems that the strains studied with genomics throughout the years have been isolated from the very same hosts these bacteria can cause disease in highly relevant crops. However, there are several reports of *Clavibacter* strains being isolated from a variety of plant sources like plum (Janisiewicz et al., 2013), coffee (Vega et al., 2005), poplar (Ulrich et al., 2008) and even wild plants (Ding et al., 2011; Zinniel et al., 2002). Moreover, several of the strains included in our analysis were isolated from sources different to those commonly associated with these pathogenic species. Hence, *Clavibacter* bacteria are not restricted to agricultural systems, and they can thrive in wild plant populations, as evidenced by the isolation of RA1B from a wild variety of tomato.

Unexpectedly, plants belonging to the *Poaceae* and *Solanaceae* plant families are the most frequent hosts of *Clavibacter* strains. These plant families are not closely related as shown by previous phylogenetic reconstructions (Leebens-Mack et al., 2019; Särkinen et al., 2013; Soreng et al., 2017), which clearly contrast with the close association between several *Clavibacter* strains obtained from such diverse hosts. The occurrence of such relationships throughout the phylogeny, at subclade and clade levels, and the incongruence between the *Clavibacter* genus phylogeny and those of their hosts, indicates lack of plant-bacteria co-evolution. Host shift is seen as a mechanism used by bacteria for long-term survival, as it allows pathogens to evolve and diversify through radiation and speciation (Langridge et al., 2015; Thines, 2019). Since host-shifting seems to be common trend in the Clavibacter genus, as shown herein, its very likely that the known pathogenic species appeared because of such mechanism after host shifting events. Even though *C. michiganensis* is the *Clavibacter* species typically associated with tomato (*S. lycopersicum*) the existence of several tomato isolates not belonging to this species and spread throughout the genus phylogeny is indicative of this phenomenon. As several conditions are required to perform a successful host shift, this observation suggests pre-adaptation as well as the occurrence of a series of events allowing these bacteria to overcome ecological barriers and colonize tomato plants, which could be related to current production practices (Anzalone et al., 2022; Hao et al., 2019; Ristaino et al., 2021; Wyngaard & Kissinger, 2022) or the impact of domestication and breeding on the plant’s physiology and its microbiome (Carrillo et al., 2019; Jaiswal et al., 2020; Soldan et al., 2021). The existence of *C. michiganensis* strains isolated from grasses at the base of this subclade (**Fig. 1**) supports the hypothesis of *C. michiganensis*’ origin outside of tomato. This is further supported by our inability to isolate *C. michiganensis* from wild tomato plants, both in Mexico and in Chile (M. Valenzuela, unpublished).

Based on in-depth genomics analyses we were able to identify the mark of key signatures of adaptation in the genetic composition of *C. michiganensis*. Our pangenomic analysis highlights that the acquisition of genes represents a key turning point during the evolution of *C. michiganensis* as a tomato pathogen, providing experimentally testable hypotheses. For instance, mutants lacking the PAI, whose genes were identified by our analysis, have been shown to have reduced virulence (Gartemann et al., 2008). Interestingly, *C. michiganensis* strain VQ143, which lacks most of the PAI genes we identified, has shown a low degree of virulence when tested *in planta* (Valenzuela et al., 2021). Identification of conserved gene families in *C. michiganensis* encoding for carbohydrate and nitrogen compounds transporter proteins, suggests that nutrient acquisition strategies were relevant for the adaptation of this bacteria to the tomato xylem environment. Given that nutrient acquiring adaptations are key for bacterial pathogens, and endophytes alike, to thrive in the poor nutrient environment provided by the xylem, nutrients are not only used for metabolism but also as signals that can trigger environmental-driven specific responses (De La Fuente et al., 2022).

The occurrence of RiPP encoding BGCs as a distinctive feature of *C. michiganensis* and related species suggests that dealing with bacterial competitors was an important adaptation to the tomato environment. It has been reported that, as in the case of michiganin, the molecules produced by these BGCs have the capacity to inhibit the growth of closely related bacteria, including other *Clavibacter* species (Holtsmark et al., 2006). However, the compounds produced by RiPP BGCs could have other roles different than antibiosis, as their ability to mediate intra-specific and host-bacteria interactions is well-acknowledged (Li & Rebuffat, 2020). The most parsimonious mechanism for spreading of these BGCs seems horizontal gene transfer, again speaking out of the contribution of RiPPs towards adaptation and evolution of pathogenic lifestyle.

In summary, our long-term and in-depth evolutionary genomics analyses – including new data doubling the number of *C. michiganensis* genome sequences available– solves a long-standing mystery about the origin and unpredictable pathogenic behavior of *C. michiganensis* causing bacterial canker. A better understanding of the evolutionary history of this seed-borne pathogen, explained by a host shift from a grass to tomato, lightens up several possibilities for its control and diagnostic, and for avoiding the occurrence of similar scenarios involving other *Clavibacter* species (*e.g. C. nebraskensis*; Osdaghi et al., 2023; Webster et al., 2019). On one hand, we anticipate that experimental validation of the candidate genes we have identified here will provide a complete picture of the pathogenicity of this endophyte. This in turn will make appearance of the disease in crops more predictable, traceable, and ultimately, controllable. On the other hand, it also provides lessons about pitfalls during plant genetic breeding, which represents the most likely place and moment for a host shift involving a seed-borne endophyte to occur. In this respect, fighting the enemy from within, *e.g.* by incorporating plant rewilding (Chen et al., 2017) and/or microbiome engineering (Compant et al., 2019; Noman et al., 2021) strategies, seems a more sustainable solution than treating this phytopathogen as an opportunist with the high costs associated with a ‘search and destroy’ strategy based on disinfectants, so-called certified seeds and/or agrochemicals.

## Data availability

Genomes from *Clavibacter* isolates obtained in Mexico, Uruguay and the Netherlands used for this research will be released as part of the BioProject PRJNA996097.

## Attributions

The following plant pictures in Figure 1 were obtained from these sources: *S. lycopersicum* from “Greenhouse Israel_IMG_3118.JPG” by Eddau, which is licensed under <u>CC BY-SA 3.0</u>; *D. glomerata* from “DactylisGlomerataIreland.JPG” by Notafly2, which is licensed under <u>CC BY-SA 3.0</u>; *E. repens* from “Elymus repens (3738612411).jpg” by Matt Lavin, which is licensed under <u>CC BY-SA 2.0</u>; *Agrostis sp.* from “Gewoon struisgras Agrostis tenuis.jpg” by Rasbak, which is licensed under <u>CC BY-SA 3.0</u>; *S. tuberosum* from “Starr 020701-0018 Solanum tuberosum.jpg” by Forest & Kim Starr, which is licensed under <u>CC BY 3.0</u>; *Z. mays* from “ZeaMays.jpg” by Christian Fischer, which is licensed under <u>CC BY-SA 3.0</u>; *M. sativa* from “Medicago sativa Alfalfa ლურჯი იონჯა.JPG” by Lazaregagnidze, which is licensed under <u>CC BY-SA 4.0</u>; *M. truncatula* from “Medicago truncatula habit1 Denman – 30067303043.jpg” by Macleay Grass Man, which is licensed under <u>CC BY 2.0</u>; *P. virgatum* from “Panicum virgatum Shenandoah 13zz.jpg” by David J. Stang, which is licensed under <u>CC BY-SA 4.0</u>; *P. vulgaris* from “Snijboon peulen Phaseolus vulgaris.jpg” by Rasbak; which is licensed under <u>CC BY-SA 3.0</u>; *S. racemosa* from “Red-berried Elder (Sambucus racemosa) – Oslo, Norway 2020-08-03 (02).jpg” by Ryan Hodnett; which is licensed under <u>CC BY-SA 4.0</u>; *T. aestivum* from “Starr-160415-0766-Triticum aestivum-home grown-Hawea Pl Olinda-Maui (26352044963).jpg” by Forest & Kim Starr, which is licensed under <u>CC BY 3.0 US</u> ; *Tulipa sp.* from “Tulipa Golden Apeldoorn.jpg” by elPadawan, which is licensed under <u>CC BY-SA 2.0</u>; *C. annum* from “Sibirischer Hauspaprika mit noch grünen Chilis.jpg” by Singlespeedfahrer, which is licensed under <u>CC0 1.0</u> and *L. perenne* from “Lolium perenne Engels raaigras.jpg” by Rasbak, which is licensed under <u>CC BY-SA 3.0</u>. All images were solely cropped to the same size, no other modifications were performed.

## Acknowledgements

This work was funded by Koppert Mexico SA de CV, Endogenomiks SAPI and Conacyt, Mexico (CB-2016 285746). FBG gratefully acknowledges support from Koppert BV, the Netherlands, during a sabbatical stay; and AYO from Conacyt, Mexico (748806) and the Institute of Biology, Leiden University, to complete PhD research. We are also grateful to Jose Luis Steffani Vallejo, Leslie Martinez Silva and Martha Angélica Martínez Tovar (Mexico) for their technical support in the greenhouse assays, and to Roberto Aviña Carlin and Alvaro Nieto for field work support.

