## Supplemental Material- Figures and tables for "A host shift as the origin of tomato bacterial canker caused by *Clavibacter michiganensis*"

### Supplementary Figures

**Supplementary figure 1**. Phylogeny of the genus *Clavibacter*

**Supplementary figure 2.** Average Nucleotide Identity analysis of *Clavibacter* genus

### Supplementary Tables

**Supplementary table S1**: List of isolates obtained from wild tomato variety populations.

**Supplementary table S2**: ANI values from pairwise comparison of the Clavibacter genus DB.

**Supplementary table S3**: *Clavibacter* genus genome database information.

**Supplementary table 4**: Gene families identified by the pangenomic and enrichment analysis

### Supplementary figures

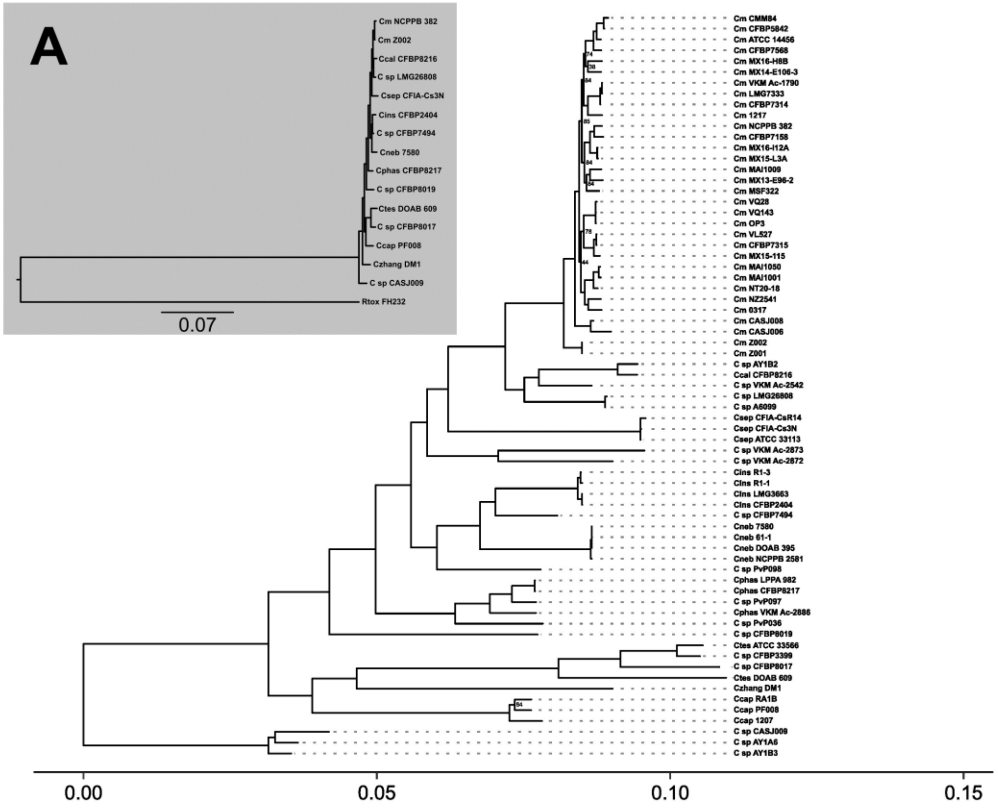


**Supplementary figure 1. Phylogeny of the genus *Clavibacter*.** Family-level reference phylogeny with *Rathayibacter toxicus* FH232 (Rtox FH232) in Subpanel A was used to order the *Clavibacter* genus phylogeny comprised of 69 genomes. Only bootstrap values <90 are displayed. *Cm, C. michiganensis; C sp., Clavibacter* sp.; *Ccal, C. californiensis*; *Csep*, *C. sepedonicus*; *Cneb*, *C. nebraskensis*; *Cins, C. insidiosus*; *Cphas*, *C. phaseoli*; *Czhang, C. zhanzhiyongii*; *Ccap, C. capsici*; *Ctes, C. tessellarius*.

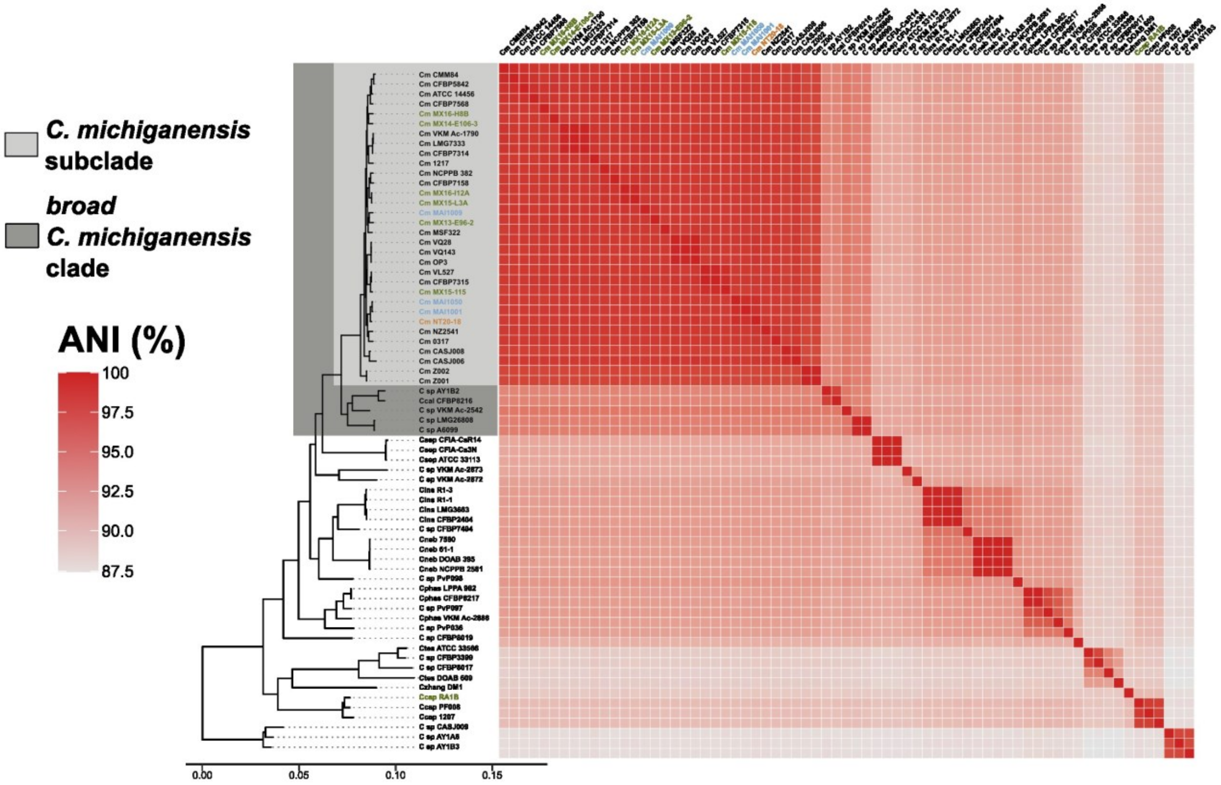


**Supplementary figure 2. Average Nucleotide Identity analysis of *Clavibacter* genus.** Heatmap showing an ANI pairwise comparison using PyANI. Rows and columns are ordered according to the genus phylogeny. Names colored other than black indicate the genomes obtained by us and published as part of this study. The colors in the names correspond to the strains' place of origin: orange - the Netherlands, blue – Uruguay, and green - Mexico.

### Supplementary tables

**Supplementary table S1.** List of isolates obtained from wild tomato variety populations.

| Isolate number | Plant sample number | Location of plan sample (Population) | Genus taxonomic identity (16SrRNA sequence) |
|---|---|---|---|
| **1** | 1 | Guanajuato | Curtobacterium |
| **2** | 1 | Guanajuato | Microbacterium |
| **3** | 1 | Guanajuato | Curtobacterium |
| **4** | 2 | Jalisco | Microbacterium |
| **5** | 3 | Jalisco | Agrococcus |
| **6** | 3 | Jalisco | Microbacterium |
| **7** | 3 | Jalisco | Microbacterium |
| **8** | 3 | Jalisco | Microbacterium |
| **9** | 3 | Jalisco | Agrococcus |
| **10** | 3 | Jalisco | Kocuria |
| **11** | 4 | Jalisco | Microbacterium |
| **12** | 4 | Jalisco | Microbacterium |
| **13** | 4 | Jalisco | Curtobacterium |
| **14** | 4 | Jalisco | Curtobacterium |
| **15** | 4 | Jalisco | Microbacterium |
| **16** | 5 | Jalisco | Curtobacterium |
| **17** | 5 | Jalisco | Microbacterium |
| **18** | 5 | Jalisco | Curtobacterium |
| **19** | 5 | Jalisco | Frigobacterium |
| **20** | 5 | Jalisco | Agrococcus |
| **21** | 6 | Guanajuato | Curtobacterium |
| **22** | 6 | Guanajuato | Curtobacterium |
| **23** | 6 | Guanajuato | Curtobacterium |
| **24** | 6 | Guanajuato | Leucobacter |
| **25** | 6 | Guanajuato | Arthrobacter |
| **26** | 6 | Guanajuato | Microbacterium |
| **27** | 7 | Guanajuato | Curtobacterium |
| **28** | 7 | Guanajuato | Arthrobacter |
| **29** | 7 | Guanajuato | Arthrobacter |
| **30** | 7 | Guanajuato | Microbacterium |
| **31** | 7 | Guanajuato | Arsenicoccus |
| **32** | 7 | Guanajuato | Curtobacterium |
| **33** | 7 | Guanajuato | Labedella |
| **34** | 8 | Guanajuato | Curtobacterium |

| | | | |
|---|---|---|---|
| **35** | 8 | Guanajuato | Curtobacterium |
| **36** | 8 | Guanajuato | Arthrobacter |
| **37** | 8 | Guanajuato | Brevibacterium |
| **38** | 8 | Guanajuato | Curtobacterium |
| **39** | 9 | Guanajuato | Microbacterium |
| **40** | 9 | Guanajuato | Microbacterium |
| **41** | 9 | Guanajuato | Microbacterium |
| **42** | 9 | Guanajuato | Cellulosimicrobium |
| **43** | 9 | Guanajuato | Microbacterium |
| **44** | 9 | Guanajuato | Okibacterium |
| **45** | 9 | Guanajuato | Microbacterium |
| **46** | 9 | Guanajuato | Microbacterium |
| **47** | 10 | Jalisco | Citricoccus |
| **48** | 10 | Jalisco | Arthrobacter |
| **49** | 10 | Jalisco | Microbacterium |
| **50** | 10 | Jalisco | Curtobacterium |
| **51** | 10 | Jalisco | Rumeliibacillus |
| **52** | 10 | Jalisco | Curtobacterium |
| **53** | 10 | Jalisco | Curtobacterium |
| **54** | 10 | Jalisco | Curtobacterium |
| **55** | 10 | Jalisco | Curtobacterium |
| **56** | 11 | Jalisco | Curtobacterium |
| **57** | 11 | Jalisco | Curtobacterium |
| **58** | 11 | Jalisco | Curtobacterium |
| **59** | 11 | Jalisco | Microbacterium |
| **60** | 11 | Jalisco | Curtobacterium |
| **61** | 11 | Jalisco | Kocuria |
| **62** | 11 | Jalisco | Brevibacterium |
| **63** | 12 | Jalisco | Curtobacterium |
| **64** | 12 | Jalisco | Microbacterium |
| **65** | 12 | Jalisco | Curtobacterium |
| **66** | 12 | Jalisco | Curtobacterium |
| **67** | 12 | Jalisco | Curtobacterium |
| **68** | 13 | Jalisco | Curtobacterium |
| **69** | 13 | Jalisco | Curtobacterium |
| **70** | 14 | Jalisco | Curtobacterium |
| **71** | 14 | Jalisco | Curtobacterium |
| **72** | 14 | Jalisco | Microbacterium |
| **73** | 14 | Jalisco | Curtobacterium |
| **74** | 15 | Jalisco | Curtobacterium |
| **75** | 15 | Jalisco | Microbacterium |
| **76** | 15 | Jalisco | Microbacterium |
| **77** | 15 | Jalisco | Kocuria |
| **78** | 16 | Jalisco | Arthrobacter |

| | | | |
|---|---|---|---|
| **79** | 16 | Jalisco | Microbacterium |
| **80** | 16 | Jalisco | Frigobacterium |
| **81** | 16 | Jalisco | Microbacterium |
| **82** | 17 | Jalisco | Curtobacterium |
| **83** | 17 | Jalisco | Curtobacterium |
| **84** | 17 | Jalisco | Curtobacterium |
| **85** | 18 | Guanajuato | Microbacterium |
| **86** | 18 | Guanajuato | Curtobacterium |
| **87** | 18 | Guanajuato | Curtobacterium |
| **88** | 18 | Guanajuato | Curtobacterium |
| **89** | 18 | Guanajuato | Plantibacter |
| **90** | 18 | Guanajuato | Curtobacterium |
| **91** | 19 | Guanajuato | Curtobacterium |
| **92** | 19 | Guanajuato | Curtobacterium |
| **93** | 19 | Guanajuato | Microbacterium |
| **94** | 19 | Guanajuato | Curtobacterium |
| **95** | 19 | Guanajuato | Curtobacterium |
| **96** | 19 | Guanajuato | Microbacterium |
| **97** | 19 | Guanajuato | Microbacterium |
| **98** | 20 | Jalisco | Curtobacterium |
| **99** | 20 | Jalisco | Curtobacterium |
| **100** | 20 | Jalisco | Curtobacterium |
| **101** | 20 | Jalisco | Microbacterium |
| **102** | 20 | Jalisco | Staphylococcus |
| **103** | 20 | Jalisco | Corynebacterium |
| **104** | 20 | Jalisco | Microbacterium |
| **105** | 20 | Jalisco | Arthrobacter |
| **106** | 20 | Jalisco | Corynebacterium |
| **107** | 20 | Jalisco | Leifsonia |
| **108** | 20 | Jalisco | Microbacterium |
| **109** | 20 | Jalisco | Arthrobacter |
| **110** | 20 | Jalisco | Microbacterium |
| **111** | 20 | Jalisco | Cellulosimicrobium |
| **112** | 20 | Jalisco | Microbacterium |
| **113** | 20 | Jalisco | Microbacterium |
| **114** | 20 | Jalisco | Microbacterium |
| **115** | 20 | Jalisco | Microbacterium |
| **116** | 20 | Jalisco | Microbacterium |
| **117** | 20 | Jalisco | Microbacterium |
| **118** | 20 | Jalisco | Arthrobacter |
| **119** | 20 | Jalisco | Microbacterium |
| **120** | 20 | Jalisco | Microbacterium |
| **121** | 20 | Jalisco | Curtobacterium |
| **122** | 21 | Jalisco | Microbacterium |

| | | | |
|---|---|---|---|
| **123** | 21 | Jalisco | Microbacterium |
| **124** | 21 | Jalisco | Curtobacterium |
| **125** | 21 | Jalisco | Curtobacterium |
| **126** | 21 | Jalisco | Microbacterium |
| **127** | 21 | Jalisco | Microbacterium |
| **128** | 21 | Jalisco | Microbacterium |
| **129** | 21 | Jalisco | Brevibacterium |
| **130** | 21 | Jalisco | Curtobacterium |
| **131** | 21 | Jalisco | Brevibacterium |
| **132** | 22 | Jalisco | Leucobacter |
| **133** | 22 | Jalisco | Brevibacterium |
| **134** | 22 | Jalisco | Brevibacterium |
| **135** | 22 | Jalisco | Curtobacterium |
| **136** | 22 | Jalisco | Microbacterium |
| **137** | 22 | Jalisco | Curtobacterium |
| **138** | 22 | Jalisco | Kocuria |
| **139** | 22 | Jalisco | Curtobacterium |
| **140** | 22 | Jalisco | Brevibacterium |
| **141** | 22 | Jalisco | Curtobacterium |
| **142** | 22 | Jalisco | Leucobacter |
| **143** | 22 | Jalisco | Brevibacterium |
| **144** | 23 | Jalisco | Leucobacter |
| **145** | 23 | Jalisco | Micrococcus |
| **146** | 23 | Jalisco | Leucobacter |
| **147** | 23 | Jalisco | Curtobacterium |
| **148** | 23 | Jalisco | Leucobacter |
| **149** | 23 | Jalisco | Microbacterium |
| **150** | 24 | Jalisco | Microbacterium |
| **151** | 24 | Jalisco | Curtobacterium |
| **152** | 24 | Jalisco | Georgenia |
| **153** | 24 | Jalisco | Microbacterium |
| **154** | 24 | Jalisco | Staphylococcus |
| **155** | 24 | Jalisco | Microbacterium |
| **156** | 24 | Jalisco | Curtobacterium |
| **157** | 24 | Jalisco | Microbacterium |
| **158** | 24 | Jalisco | Microbacterium |
| **159** | 24 | Jalisco | Microbacterium |
| **160** | 24 | Jalisco | Microbacterium |
| **161** | 24 | Jalisco | Frigoribacterium |
| **162** | 24 | Jalisco | Leucobacter |
| **163** | 24 | Jalisco | Microbacterium |
| **164** | 24 | Jalisco | Curtobacterium |
| **165** | 24 | Jalisco | Bacillus |
| **166** | 24 | Jalisco | Rhodococcus |

| | | | |
|---|---|---|---|
| **167** | 24 | Jalisco | Bacillus |
| **168** | 24 | Jalisco | Agrococcus |
| **169** | 24 | Jalisco | Frigoribacterium |
| **170** | 24 | Jalisco | Pseudarthrobacter |
| **171** | 24 | Jalisco | Rhodococcus |
| **172** | 24 | Jalisco | Staphylococcus |
| **173** | 24 | Jalisco | Kocuria |
| **174** | 25 | Guanajuato | Microbacterium |
| **175** | 25 | Guanajuato | Kocuria |
| **176** | 25 | Guanajuato | Microbacterium |
| **177** | 25 | Guanajuato | Agrococcus |
| **178** | 26 | Guanajuato | Curtobacterium |
| **179** | 26 | Guanajuato | Curtobacterium |
| **180** | 26 | Guanajuato | Curtobacterium |
| **181** | 26 | Guanajuato | Pseudarthrobacter |
| **182** | 26 | Guanajuato | Microbacterium |
| **183** | 26 | Guanajuato | Arthrobacter |
| **184** | 26 | Guanajuato | Arthrobacter |
| **185** | 26 | Guanajuato | Microbacterium |
| **186** | 26 | Guanajuato | Curtobacterium |
| **187** | 26 | Guanajuato | Sanguibacter |
| **188** | 26 | Guanajuato | Labedella |
| **189** | 26 | Guanajuato | Curtobacterium |
| **190** | 26 | Guanajuato | Curtobacterium |
| **191** | 26 | Guanajuato | Microbacterium |
| **192** | 26 | Guanajuato | Curtobacterium |
| **193** | 26 | Guanajuato | Curtobacterium |
| **194** | 26 | Guanajuato | Microbacterium |
| **195** | 26 | Guanajuato | Curtobacterium |
| **196** | 26 | Guanajuato | Curtobacterium |
| **197** | 26 | Guanajuato | Microbacterium |
| **198** | 26 | Guanajuato | Curtobacterium |
| **199** | 26 | Guanajuato | Paenarthrobacter |
| **200** | 26 | Guanajuato | Microbacterium |
| **201** | 26 | Guanajuato | Microbacterium |
| **202** | 26 | Guanajuato | Curtobacterium |
| **203** | 26 | Guanajuato | Curtobacterium |
| **204** | 26 | Guanajuato | Curtobacterium |
| **205** | 26 | Guanajuato | Microbacterium |
| **206** | 26 | Guanajuato | Microbacterium |
| **207** | 26 | Guanajuato | Curtobacterium |
| **208** | 27 | Guanajuato | Microbacterium |
| **209** | 27 | Guanajuato | Clavibacter |
| **210** | 27 | Guanajuato | Microbacterium |

| | | | |
|---|---|---|---|
| **211** | 28 | Guanajuato | Bacillus |
| **212** | 28 | Guanajuato | Microbacterium |
| **213** | 28 | Guanajuato | Arthrobacter |
| **214** | 28 | Guanajuato | Microbacterium |
| **215** | 28 | Guanajuato | Microbacterium |
| **216** | 28 | Guanajuato | Curtobacterium |
| **217** | 28 | Guanajuato | Curtobacterium |
| **218** | 28 | Guanajuato | Arthrobacter |
| **219** | 28 | Guanajuato | Microbacterium |
| **220** | 28 | Guanajuato | Arthrobacter |
| **221** | 28 | Guanajuato | Arthrobacter |
| **222** | 28 | Guanajuato | Microbacterium |

**Supplementary table S2**: ANI values from pairwise comparison of the Clavibacter genus DB. Cells are colored in a color gradient from white to red according to the lowest (0.874) and highest (1.00) ANI values. *Cm, C. michiganensis; C sp., Clavibacter* sp.; *Ccal, C. californiensis*; *Csep*, *C. sepedonicus*; *Cneb*, *C. nebraskensis*; *Cins, C. insidiosus*; *Cphas*, *C. phaseoli*; *Czhang, C. zhanzhiyongii*; *Ccap, C. capsici*; *Ctes, C. tessellarius*.

| | Cm CFBP5842 | Cm CMM84 | Cm ATCC 14456 | Cm CFBP7568 | Cm MX14-E106-3 | Cm MX16-H8B | Cm LMG7333 | Cm VKM Ac-1790 | Cm CFBP7314 | Cm 1217 | Cm MX15-L3A | Cm MX16-I12A | Cm CFBP7158 | Cm NCPPB 382 | Cm MX13-E96-2 | Cm MAI1009 | Cm MSF322 | Cm CFBP7315 | Cm VL527 | Cm MX15-115 | Cm VQ143 |
|---|---|---|---|---|---|---|---|---|---|---|---|---|---|---|---|---|---|---|---|---|---|
| Cm CFBP5842 | 1.000 | 0.998 | 0.996 | 0.994 | 0.992 | 0.991 | 0.991 | 0.992 | 0.992 | 0.991 | 0.991 | 0.991 | 0.991 | 0.991 | 0.991 | 0.992 | 0.991 | 0.991 | 0.991 | 0.991 | 0.990 |
| Cm CMM84 | 0.998 | 1.000 | 0.995 | 0.993 | 0.992 | 0.991 | 0.993 | 0.993 | 0.993 | 0.992 | 0.991 | 0.991 | 0.991 | 0.991 | 0.991 | 0.992 | 0.990 | 0.991 | 0.990 | 0.990 | 0.990 |
| Cm ATCC 14456 | 0.996 | 0.995 | 1.000 | 0.995 | 0.992 | 0.993 | 0.992 | 0.992 | 0.992 | 0.992 | 0.992 | 0.992 | 0.992 | 0.992 | 0.992 | 0.993 | 0.991 | 0.992 | 0.992 | 0.991 | 0.991 |
| Cm CFBP7568 | 0.994 | 0.993 | 0.995 | 1.000 | 0.992 | 0.992 | 0.991 | 0.991 | 0.992 | 0.991 | 0.991 | 0.991 | 0.992 | 0.991 | 0.991 | 0.992 | 0.991 | 0.991 | 0.991 | 0.991 | 0.990 |
| Cm MX14-E106-3 | 0.991 | 0.991 | 0.991 | 0.991 | 1.000 | 0.991 | 0.990 | 0.990 | 0.990 | 0.991 | 0.990 | 0.990 | 0.990 | 0.991 | 0.990 | 0.991 | 0.989 | 0.990 | 0.990 | 0.990 | 0.989 |
| Cm MX16-H8B | 0.991 | 0.991 | 0.991 | 0.991 | 0.991 | 1.000 | 0.990 | 0.991 | 0.991 | 0.991 | 0.990 | 0.991 | 0.991 | 0.990 | 0.991 | 0.990 | 0.990 | 0.991 | 0.989 | 0.990 | 0.990 |
| Cm LMG7333 | 0.991 | 0.993 | 0.991 | 0.991 | 0.991 | 0.991 | 1.000 | 1.000 | 1.000 | 0.993 | 0.991 | 0.991 | 0.992 | 0.991 | 0.992 | 0.992 | 0.990 | 0.991 | 0.991 | 0.990 | 0.991 |
| Cm VKM Ac-1790 | 0.992 | 0.993 | 0.992 | 0.991 | 0.991 | 0.992 | 1.000 | 1.000 | 1.000 | 0.993 | 0.992 | 0.992 | 0.992 | 0.992 | 0.992 | 0.992 | 0.991 | 0.992 | 0.992 | 0.991 | 0.991 |
| Cm CFBP7314 | 0.992 | 0.993 | 0.992 | 0.991 | 0.991 | 0.992 | 1.000 | 1.000 | 1.000 | 0.993 | 0.992 | 0.992 | 0.992 | 0.992 | 0.992 | 0.992 | 0.991 | 0.991 | 0.992 | 0.991 | 0.991 |
| Cm 1217 | 0.991 | 0.991 | 0.991 | 0.991 | 0.992 | 0.991 | 0.993 | 0.992 | 0.992 | 1.000 | 0.991 | 0.991 | 0.991 | 0.990 | 0.991 | 0.992 | 0.990 | 0.991 | 0.990 | 0.990 | 0.990 |
| Cm MX15-L3A | 0.990 | 0.991 | 0.991 | 0.991 | 0.991 | 0.991 | 0.991 | 0.991 | 0.991 | 0.991 | 1.000 | 0.999 | 0.994 | 0.994 | 0.992 | 0.992 | 0.991 | 0.991 | 0.991 | 0.991 | 0.990 |
| Cm MX16-I12A | 0.991 | 0.991 | 0.991 | 0.991 | 0.991 | 0.991 | 0.991 | 0.992 | 0.991 | 0.991 | 0.999 | 1.000 | 0.994 | 0.994 | 0.992 | 0.991 | 0.991 | 0.991 | 0.990 | 0.990 | 0.990 |
| Cm CFBP7158 | 0.991 | 0.991 | 0.992 | 0.991 | 0.991 | 0.991 | 0.992 | 0.992 | 0.992 | 0.992 | 0.995 | 0.995 | 1.000 | 0.995 | 0.992 | 0.992 | 0.991 | 0.991 | 0.991 | 0.991 | 0.990 |
| Cm NCPPB 382 | 0.991 | 0.991 | 0.991 | 0.991 | 0.992 | 0.991 | 0.991 | 0.992 | 0.992 | 0.991 | 0.994 | 0.994 | 0.995 | 1.000 | 0.992 | 0.992 | 0.991 | 0.991 | 0.991 | 0.991 | 0.990 |
| Cm MX13-E96-2 | 0.991 | 0.991 | 0.991 | 0.991 | 0.991 | 0.992 | 0.991 | 0.992 | 0.991 | 0.991 | 0.992 | 0.992 | 0.991 | 0.991 | 1.000 | 0.992 | 0.990 | 0.991 | 0.990 | 0.990 | 0.990 |
| Cm MAI1009 | 0.992 | 0.992 | 0.992 | 0.991 | 0.992 | 0.991 | 0.992 | 0.992 | 0.992 | 0.992 | 0.992 | 0.992 | 0.992 | 0.992 | 0.992 | 1.000 | 0.991 | 0.991 | 0.991 | 0.990 | 0.990 |
| Cm MSF322 | 0.991 | 0.989 | 0.991 | 0.990 | 0.991 | 0.990 | 0.990 | 0.991 | 0.991 | 0.990 | 0.991 | 0.991 | 0.991 | 0.990 | 0.991 | 0.991 | 1.000 | 0.991 | 0.992 | 0.990 | 0.991 |
| Cm CFBP7315 | 0.992 | 0.991 | 0.992 | 0.992 | 0.991 | 0.992 | 0.992 | 0.992 | 0.992 | 0.992 | 0.992 | 0.991 | 0.991 | 0.992 | 0.992 | 0.992 | 0.992 | 1.000 | 1.000 | 0.998 | 0.992 |
| Cm VL527 | 0.991 | 0.990 | 0.991 | 0.991 | 0.991 | 0.991 | 0.991 | 0.991 | 0.991 | 0.990 | 0.991 | 0.991 | 0.991 | 0.991 | 0.991 | 0.991 | 0.992 | 0.999 | 1.000 | 0.996 | 0.991 |
| Cm MX15-115 | 0.990 | 0.989 | 0.991 | 0.990 | 0.990 | 0.990 | 0.990 | 0.991 | 0.991 | 0.990 | 0.990 | 0.990 | 0.990 | 0.990 | 0.990 | 0.990 | 0.990 | 0.997 | 0.996 | 1.000 | 0.990 |
| Cm VQ143 | 0.990 | 0.991 | 0.991 | 0.990 | 0.990 | 0.990 | 0.992 | 0.991 | 0.991 | 0.991 | 0.991 | 0.990 | 0.991 | 0.990 | 0.991 | 0.991 | 0.991 | 0.992 | 0.991 | 0.991 | 1.000 |
| Cm VQ28 | 0.991 | 0.991 | 0.991 | 0.991 | 0.990 | 0.991 | 0.991 | 0.991 | 0.991 | 0.991 | 0.991 | 0.991 | 0.991 | 0.991 | 0.991 | 0.991 | 0.992 | 0.992 | 0.992 | 0.992 | 0.999 |
| Cm OP3 | 0.991 | 0.991 | 0.991 | 0.990 | 0.990 | 0.990 | 0.991 | 0.991 | 0.991 | 0.991 | 0.991 | 0.990 | 0.991 | 0.990 | 0.991 | 0.991 | 0.991 | 0.992 | 0.991 | 0.991 | 0.999 |
| Cm MAI1001 | 0.991 | 0.990 | 0.991 | 0.990 | 0.991 | 0.990 | 0.991 | 0.991 | 0.991 | 0.991 | 0.991 | 0.991 | 0.991 | 0.991 | 0.991 | 0.991 | 0.991 | 0.991 | 0.991 | 0.990 | 0.991 |
| Cm MAI1050 | 0.990 | 0.990 | 0.991 | 0.990 | 0.990 | 0.989 | 0.990 | 0.991 | 0.991 | 0.990 | 0.990 | 0.990 | 0.990 | 0.990 | 0.990 | 0.990 | 0.989 | 0.991 | 0.990 | 0.990 | 0.990 |
| Cm NT20-18 | 0.991 | 0.990 | 0.991 | 0.990 | 0.991 | 0.990 | 0.991 | 0.991 | 0.991 | 0.991 | 0.991 | 0.990 | 0.991 | 0.990 | 0.991 | 0.991 | 0.991 | 0.991 | 0.991 | 0.991 | 0.992 |
| Cm 0317 | 0.990 | 0.990 | 0.991 | 0.990 | 0.990 | 0.990 | 0.991 | 0.991 | 0.991 | 0.991 | 0.991 | 0.991 | 0.991 | 0.991 | 0.991 | 0.991 | 0.991 | 0.992 | 0.991 | 0.991 | 0.990 |
| Cm NZ2541 | 0.990 | 0.990 | 0.991 | 0.990 | 0.990 | 0.990 | 0.991 | 0.991 | 0.991 | 0.991 | 0.991 | 0.991 | 0.991 | 0.991 | 0.991 | 0.991 | 0.991 | 0.991 | 0.991 | 0.991 | 0.991 |
| Cm CASJ006 | 0.991 | 0.990 | 0.991 | 0.990 | 0.990 | 0.990 | 0.991 | 0.991 | 0.991 | 0.990 | 0.991 | 0.991 | 0.991 | 0.991 | 0.991 | 0.991 | 0.991 | 0.991 | 0.991 | 0.991 | 0.991 |
| Cm CASJ008 | 0.988 | 0.987 | 0.988 | 0.987 | 0.987 | 0.988 | 0.988 | 0.988 | 0.988 | 0.989 | 0.988 | 0.988 | 0.988 | 0.988 | 0.988 | 0.988 | 0.988 | 0.988 | 0.988 | 0.988 | 0.988 |
| Cm Z001 | 0.986 | 0.985 | 0.986 | 0.985 | 0.985 | 0.985 | 0.985 | 0.985 | 0.986 | 0.986 | 0.985 | 0.986 | 0.986 | 0.986 | 0.985 | 0.985 | 0.985 | 0.986 | 0.986 | 0.985 | 0.986 |
| Cm Z002 | 0.985 | 0.985 | 0.986 | 0.985 | 0.985 | 0.985 | 0.985 | 0.985 | 0.985 | 0.985 | 0.985 | 0.985 | 0.986 | 0.985 | 0.985 | 0.984 | 0.985 | 0.985 | 0.985 | 0.985 | 0.985 |
| Ccal CFBP8216 | 0.944 | 0.943 | 0.944 | 0.943 | 0.944 | 0.944 | 0.943 | 0.944 | 0.944 | 0.944 | 0.944 | 0.944 | 0.944 | 0.943 | 0.943 | 0.943 | 0.943 | 0.944 | 0.943 | 0.944 | 0.943 |
| C sp AY1B2 | 0.943 | 0.942 | 0.943 | 0.943 | 0.943 | 0.942 | 0.942 | 0.943 | 0.943 | 0.943 | 0.943 | 0.943 | 0.943 | 0.942 | 0.942 | 0.942 | 0.942 | 0.943 | 0.942 | 0.942 | 0.942 |
| C sp VKM Ac-2542 | 0.952 | 0.952 | 0.952 | 0.952 | 0.952 | 0.952 | 0.952 | 0.952 | 0.952 | 0.953 | 0.952 | 0.952 | 0.952 | 0.952 | 0.952 | 0.952 | 0.952 | 0.952 | 0.952 | 0.952 | 0.952 |
| C sp A6099 | 0.948 | 0.947 | 0.948 | 0.947 | 0.948 | 0.948 | 0.947 | 0.948 | 0.948 | 0.948 | 0.947 | 0.947 | 0.947 | 0.947 | 0.947 | 0.947 | 0.947 | 0.948 | 0.947 | 0.947 | 0.947 |
| C sp LMG 26808 | 0.947 | 0.946 | 0.947 | 0.947 | 0.947 | 0.947 | 0.946 | 0.947 | 0.947 | 0.948 | 0.947 | 0.947 | 0.947 | 0.946 | 0.946 | 0.946 | 0.946 | 0.947 | 0.946 | 0.947 | 0.946 |
| Csep CFIA-Cs3N | 0.919 | 0.918 | 0.918 | 0.918 | 0.918 | 0.918 | 0.918 | 0.918 | 0.918 | 0.919 | 0.919 | 0.918 | 0.918 | 0.917 | 0.918 | 0.917 | 0.918 | 0.918 | 0.917 | 0.918 | 0.919 |
| Csep CFIA-CsR14 | 0.918 | 0.917 | 0.918 | 0.917 | 0.918 | 0.918 | 0.917 | 0.917 | 0.918 | 0.919 | 0.918 | 0.918 | 0.918 | 0.917 | 0.918 | 0.917 | 0.917 | 0.917 | 0.917 | 0.918 | 0.919 |
| Csep ATCC33113 | 0.919 | 0.918 | 0.918 | 0.918 | 0.918 | 0.918 | 0.918 | 0.918 | 0.918 | 0.920 | 0.919 | 0.919 | 0.918 | 0.918 | 0.918 | 0.918 | 0.918 | 0.918 | 0.918 | 0.918 | 0.919 |
| C sp VKM Ac-2872 | 0.924 | 0.922 | 0.923 | 0.923 | 0.923 | 0.922 | 0.922 | 0.923 | 0.923 | 0.924 | 0.923 | 0.923 | 0.923 | 0.923 | 0.923 | 0.923 | 0.923 | 0.923 | 0.923 | 0.923 | 0.923 |
| C sp VKM Ac-2873 | 0.919 | 0.918 | 0.919 | 0.918 | 0.919 | 0.919 | 0.918 | 0.918 | 0.918 | 0.920 | 0.919 | 0.919 | 0.918 | 0.918 | 0.919 | 0.918 | 0.918 | 0.919 | 0.918 | 0.919 | 0.919 |
| Cins CFBP2404 | 0.927 | 0.926 | 0.927 | 0.926 | 0.926 | 0.926 | 0.926 | 0.926 | 0.926 | 0.928 | 0.927 | 0.927 | 0.927 | 0.927 | 0.927 | 0.926 | 0.926 | 0.927 | 0.926 | 0.927 | 0.927 |
| Cins LMG 3663 | 0.927 | 0.926 | 0.926 | 0.925 | 0.926 | 0.926 | 0.926 | 0.926 | 0.926 | 0.928 | 0.927 | 0.927 | 0.926 | 0.927 | 0.927 | 0.926 | 0.925 | 0.926 | 0.926 | 0.926 | 0.927 |
| Cins R1-1 | 0.926 | 0.926 | 0.926 | 0.925 | 0.926 | 0.926 | 0.926 | 0.926 | 0.926 | 0.928 | 0.927 | 0.927 | 0.926 | 0.926 | 0.926 | 0.926 | 0.925 | 0.926 | 0.925 | 0.926 | 0.926 |
| Cins R1-3 | 0.927 | 0.926 | 0.926 | 0.926 | 0.927 | 0.926 | 0.926 | 0.926 | 0.926 | 0.928 | 0.927 | 0.927 | 0.926 | 0.926 | 0.927 | 0.926 | 0.925 | 0.926 | 0.926 | 0.926 | 0.927 |
| C sp CFBP7494 | 0.929 | 0.929 | 0.928 | 0.928 | 0.929 | 0.929 | 0.929 | 0.929 | 0.929 | 0.931 | 0.929 | 0.929 | 0.928 | 0.929 | 0.929 | 0.929 | 0.929 | 0.929 | 0.929 | 0.929 | 0.929 |
| Cneb 61-1 | 0.926 | 0.925 | 0.925 | 0.925 | 0.925 | 0.926 | 0.925 | 0.925 | 0.925 | 0.927 | 0.926 | 0.926 | 0.925 | 0.925 | 0.926 | 0.925 | 0.925 | 0.925 | 0.925 | 0.926 | 0.925 |
| Cneb 7580 | 0.926 | 0.925 | 0.926 | 0.925 | 0.925 | 0.926 | 0.925 | 0.925 | 0.925 | 0.927 | 0.926 | 0.926 | 0.925 | 0.925 | 0.926 | 0.925 | 0.925 | 0.926 | 0.925 | 0.926 | 0.925 |
| Cneb DOAB 395 | 0.926 | 0.925 | 0.925 | 0.925 | 0.925 | 0.926 | 0.925 | 0.925 | 0.925 | 0.928 | 0.926 | 0.926 | 0.925 | 0.925 | 0.926 | 0.925 | 0.925 | 0.926 | 0.926 | 0.926 | 0.925 |
| Cneb NCPPB 2581 | 0.926 | 0.925 | 0.925 | 0.924 | 0.925 | 0.925 | 0.925 | 0.925 | 0.925 | 0.927 | 0.926 | 0.925 | 0.925 | 0.925 | 0.925 | 0.925 | 0.925 | 0.925 | 0.925 | 0.925 | 0.925 |
| C sp PvP098 | 0.930 | 0.930 | 0.930 | 0.930 | 0.930 | 0.930 | 0.930 | 0.930 | 0.930 | 0.931 | 0.930 | 0.930 | 0.930 | 0.930 | 0.930 | 0.930 | 0.930 | 0.930 | 0.930 | 0.930 | 0.930 |
| Cchil CFBP8217 | 0.926 | 0.925 | 0.925 | 0.925 | 0.926 | 0.925 | 0.925 | 0.925 | 0.925 | 0.928 | 0.925 | 0.926 | 0.925 | 0.925 | 0.926 | 0.925 | 0.925 | 0.925 | 0.925 | 0.926 | 0.925 |
| Cphas LPPA 982 | 0.926 | 0.924 | 0.925 | 0.924 | 0.925 | 0.925 | 0.924 | 0.925 | 0.924 | 0.927 | 0.925 | 0.925 | 0.925 | 0.924 | 0.925 | 0.925 | 0.925 | 0.925 | 0.924 | 0.925 | 0.924 |
| C sp PvP097 | 0.926 | 0.925 | 0.925 | 0.925 | 0.925 | 0.926 | 0.925 | 0.925 | 0.925 | 0.927 | 0.926 | 0.926 | 0.925 | 0.925 | 0.926 | 0.925 | 0.925 | 0.925 | 0.925 | 0.926 | 0.925 |
| Cphas VKM Ac-2886 | 0.924 | 0.923 | 0.923 | 0.923 | 0.924 | 0.924 | 0.923 | 0.924 | 0.923 | 0.926 | 0.924 | 0.924 | 0.923 | 0.923 | 0.924 | 0.924 | 0.923 | 0.923 | 0.923 | 0.924 | 0.924 |
| C sp PvP036 | 0.923 | 0.923 | 0.922 | 0.922 | 0.923 | 0.923 | 0.922 | 0.922 | 0.922 | 0.924 | 0.923 | 0.923 | 0.922 | 0.923 | 0.923 | 0.922 | 0.922 | 0.922 | 0.922 | 0.923 | 0.922 |
| C sp CFBP8019 | 0.912 | 0.911 | 0.911 | 0.911 | 0.911 | 0.912 | 0.911 | 0.911 | 0.911 | 0.914 | 0.912 | 0.912 | 0.911 | 0.911 | 0.912 | 0.911 | 0.911 | 0.911 | 0.911 | 0.912 | 0.911 |
| C sp CFBP3399 | 0.893 | 0.892 | 0.892 | 0.892 | 0.893 | 0.893 | 0.892 | 0.892 | 0.892 | 0.896 | 0.893 | 0.893 | 0.892 | 0.892 | 0.893 | 0.892 | 0.892 | 0.892 | 0.892 | 0.893 | 0.892 |
| Ctes ATCC 33566 | 0.893 | 0.892 | 0.891 | 0.892 | 0.893 | 0.892 | 0.892 | 0.892 | 0.892 | 0.895 | 0.893 | 0.893 | 0.892 | 0.892 | 0.893 | 0.892 | 0.892 | 0.892 | 0.892 | 0.893 | 0.892 |
| C sp CFBP8017 | 0.891 | 0.889 | 0.889 | 0.889 | 0.890 | 0.890 | 0.889 | 0.889 | 0.889 | 0.893 | 0.890 | 0.890 | 0.889 | 0.890 | 0.890 | 0.889 | 0.890 | 0.889 | 0.889 | 0.890 | 0.889 |
| Ctes DOAB 609 | 0.892 | 0.891 | 0.891 | 0.890 | 0.891 | 0.891 | 0.891 | 0.891 | 0.891 | 0.894 | 0.891 | 0.891 | 0.890 | 0.891 | 0.891 | 0.891 | 0.891 | 0.891 | 0.891 | 0.892 | 0.891 |
| Czhang DM1 | 0.895 | 0.894 | 0.894 | 0.893 | 0.895 | 0.894 | 0.894 | 0.894 | 0.894 | 0.897 | 0.894 | 0.895 | 0.894 | 0.894 | 0.895 | 0.894 | 0.894 | 0.894 | 0.894 | 0.895 | 0.894 |
| Ccap PF008 | 0.903 | 0.901 | 0.902 | 0.901 | 0.902 | 0.902 | 0.901 | 0.902 | 0.902 | 0.904 | 0.902 | 0.902 | 0.902 | 0.901 | 0.901 | 0.902 | 0.901 | 0.902 | 0.901 | 0.902 | 0.902 |
| Ccap RA1B | 0.903 | 0.903 | 0.903 | 0.903 | 0.903 | 0.903 | 0.903 | 0.903 | 0.903 | 0.905 | 0.903 | 0.903 | 0.903 | 0.903 | 0.903 | 0.903 | 0.903 | 0.903 | 0.903 | 0.904 | 0.903 |
| Ccap 1207 | 0.903 | 0.902 | 0.902 | 0.902 | 0.903 | 0.902 | 0.902 | 0.902 | 0.902 | 0.904 | 0.903 | 0.902 | 0.902 | 0.902 | 0.903 | 0.902 | 0.902 | 0.903 | 0.902 | 0.903 | 0.902 |
| C sp AY1A6 | 0.881 | 0.880 | 0.880 | 0.880 | 0.881 | 0.881 | 0.880 | 0.880 | 0.880 | 0.883 | 0.881 | 0.881 | 0.880 | 0.880 | 0.881 | 0.880 | 0.880 | 0.880 | 0.880 | 0.881 | 0.880 |
| C sp CASJ009 | 0.881 | 0.880 | 0.880 | 0.880 | 0.881 | 0.881 | 0.880 | 0.880 | 0.880 | 0.883 | 0.881 | 0.881 | 0.880 | 0.880 | 0.881 | 0.880 | 0.880 | 0.880 | 0.880 | 0.882 | 0.880 |
| C sp AY1B3 | 0.881 | 0.880 | 0.880 | 0.880 | 0.881 | 0.881 | 0.880 | 0.880 | 0.880 | 0.883 | 0.881 | 0.881 | 0.880 | 0.880 | 0.881 | 0.880 | 0.881 | 0.880 | 0.880 | 0.881 | 0.880 |

| | Cm VQ28 | Cm OP3 | Cm MAI1001 | Cm MAI1050 | Cm NT20-18 | Cm 0317 | Cm NZ2541 | Cm CASJ006 | Cm CASJ008 | Cm Z001 | Cm Z002 | Ccal CFBP8216 | C sp AY1B2 | C sp VKM Ac-2542 | C sp A6099 | C sp LMG 26808 | Csep CFIA-Cs3N | Csep CFIA-CsR14 | Csep ATCC33113 | C sp VKM Ac-2872 | C sp VKM Ac-2873 | Cins CFBP2404 | Cins LMG 3663 |
|---|---|---|---|---|---|---|---|---|---|---|---|---|---|---|---|---|---|---|---|---|---|---|---|
| Cm CFBP5842 | 0.991 | 0.991 | 0.991 | 0.990 | 0.991 | 0.990 | 0.990 | 0.990 | 0.988 | 0.985 | 0.985 | 0.943 | 0.942 | 0.952 | 0.946 | 0.946 | 0.917 | 0.917 | 0.917 | 0.924 | 0.918 | 0.925 | 0.925 |
| Cm CMM84 | 0.991 | 0.991 | 0.991 | 0.990 | 0.991 | 0.990 | 0.990 | 0.990 | 0.987 | 0.985 | 0.985 | 0.943 | 0.942 | 0.952 | 0.946 | 0.946 | 0.918 | 0.918 | 0.918 | 0.923 | 0.918 | 0.926 | 0.926 |
| Cm ATCC 14456 | 0.992 | 0.992 | 0.992 | 0.991 | 0.991 | 0.991 | 0.991 | 0.991 | 0.988 | 0.986 | 0.986 | 0.944 | 0.943 | 0.953 | 0.947 | 0.946 | 0.918 | 0.918 | 0.918 | 0.923 | 0.918 | 0.926 | 0.926 |
| Cm CFBP7568 | 0.991 | 0.991 | 0.991 | 0.991 | 0.991 | 0.990 | 0.991 | 0.990 | 0.988 | 0.985 | 0.985 | 0.944 | 0.943 | 0.952 | 0.947 | 0.946 | 0.918 | 0.918 | 0.918 | 0.923 | 0.917 | 0.925 | 0.925 |
| Cm MX14-E106-3 | 0.989 | 0.989 | 0.990 | 0.989 | 0.990 | 0.989 | 0.989 | 0.989 | 0.986 | 0.984 | 0.984 | 0.943 | 0.942 | 0.951 | 0.946 | 0.946 | 0.917 | 0.917 | 0.917 | 0.922 | 0.917 | 0.925 | 0.924 |
| Cm MX16-H8B | 0.990 | 0.990 | 0.990 | 0.989 | 0.990 | 0.989 | 0.990 | 0.989 | 0.987 | 0.984 | 0.984 | 0.943 | 0.942 | 0.952 | 0.947 | 0.946 | 0.917 | 0.917 | 0.917 | 0.922 | 0.917 | 0.925 | 0.924 |
| Cm LMG7333 | 0.991 | 0.992 | 0.991 | 0.990 | 0.992 | 0.990 | 0.991 | 0.990 | 0.988 | 0.985 | 0.985 | 0.943 | 0.942 | 0.952 | 0.946 | 0.946 | 0.918 | 0.918 | 0.918 | 0.923 | 0.917 | 0.926 | 0.926 |
| Cm VKM Ac-1790 | 0.992 | 0.991 | 0.992 | 0.991 | 0.991 | 0.991 | 0.991 | 0.991 | 0.988 | 0.985 | 0.985 | 0.944 | 0.943 | 0.953 | 0.947 | 0.947 | 0.918 | 0.918 | 0.918 | 0.923 | 0.918 | 0.926 | 0.926 |
| Cm CFBP7314 | 0.992 | 0.992 | 0.991 | 0.991 | 0.992 | 0.991 | 0.991 | 0.991 | 0.988 | 0.985 | 0.985 | 0.944 | 0.943 | 0.952 | 0.947 | 0.947 | 0.918 | 0.918 | 0.918 | 0.923 | 0.917 | 0.926 | 0.926 |
| Cm 1217 | 0.991 | 0.991 | 0.991 | 0.990 | 0.991 | 0.990 | 0.990 | 0.990 | 0.988 | 0.985 | 0.985 | 0.944 | 0.943 | 0.953 | 0.947 | 0.946 | 0.918 | 0.918 | 0.918 | 0.924 | 0.919 | 0.926 | 0.926 |
| Cm MX15-L3A | 0.991 | 0.990 | 0.990 | 0.990 | 0.990 | 0.990 | 0.991 | 0.990 | 0.988 | 0.985 | 0.985 | 0.943 | 0.942 | 0.952 | 0.946 | 0.945 | 0.917 | 0.918 | 0.918 | 0.922 | 0.917 | 0.925 | 0.925 |
| Cm MX16-I12A | 0.991 | 0.991 | 0.991 | 0.990 | 0.990 | 0.990 | 0.990 | 0.990 | 0.988 | 0.985 | 0.985 | 0.943 | 0.942 | 0.952 | 0.946 | 0.946 | 0.917 | 0.917 | 0.918 | 0.922 | 0.917 | 0.926 | 0.926 |
| Cm CFBP7158 | 0.991 | 0.991 | 0.991 | 0.991 | 0.991 | 0.991 | 0.991 | 0.991 | 0.989 | 0.986 | 0.986 | 0.944 | 0.943 | 0.953 | 0.947 | 0.947 | 0.918 | 0.918 | 0.918 | 0.923 | 0.918 | 0.926 | 0.926 |
| Cm NCPPB 382 | 0.991 | 0.991 | 0.991 | 0.990 | 0.991 | 0.991 | 0.991 | 0.991 | 0.988 | 0.985 | 0.985 | 0.943 | 0.942 | 0.952 | 0.946 | 0.946 | 0.918 | 0.918 | 0.918 | 0.923 | 0.917 | 0.926 | 0.926 |
| Cm MX13-E96-2 | 0.991 | 0.991 | 0.991 | 0.990 | 0.990 | 0.990 | 0.990 | 0.990 | 0.987 | 0.984 | 0.984 | 0.943 | 0.942 | 0.952 | 0.946 | 0.945 | 0.918 | 0.918 | 0.918 | 0.923 | 0.917 | 0.926 | 0.925 |
| Cm MAI1009 | 0.991 | 0.991 | 0.991 | 0.991 | 0.991 | 0.990 | 0.990 | 0.990 | 0.988 | 0.984 | 0.984 | 0.943 | 0.942 | 0.952 | 0.947 | 0.947 | 0.918 | 0.918 | 0.918 | 0.923 | 0.918 | 0.926 | 0.926 |
| Cm MSF322 | 0.992 | 0.992 | 0.991 | 0.990 | 0.991 | 0.991 | 0.991 | 0.991 | 0.989 | 0.985 | 0.985 | 0.943 | 0.943 | 0.953 | 0.946 | 0.946 | 0.918 | 0.918 | 0.918 | 0.924 | 0.918 | 0.926 | 0.926 |
| Cm CFBP7315 | 0.992 | 0.992 | 0.992 | 0.991 | 0.992 | 0.992 | 0.991 | 0.991 | 0.989 | 0.985 | 0.985 | 0.944 | 0.943 | 0.953 | 0.947 | 0.947 | 0.917 | 0.918 | 0.917 | 0.923 | 0.917 | 0.926 | 0.926 |
| Cm VL527 | 0.992 | 0.992 | 0.991 | 0.991 | 0.991 | 0.991 | 0.991 | 0.991 | 0.988 | 0.985 | 0.985 | 0.943 | 0.943 | 0.953 | 0.946 | 0.946 | 0.917 | 0.917 | 0.917 | 0.923 | 0.918 | 0.925 | 0.925 |
| Cm MX15-115 | 0.991 | 0.990 | 0.990 | 0.989 | 0.990 | 0.990 | 0.990 | 0.990 | 0.987 | 0.984 | 0.984 | 0.943 | 0.942 | 0.952 | 0.946 | 0.946 | 0.917 | 0.917 | 0.917 | 0.923 | 0.917 | 0.925 | 0.925 |
| Cm VQ143 | 0.999 | 1.000 | 0.991 | 0.991 | 0.993 | 0.991 | 0.991 | 0.991 | 0.989 | 0.986 | 0.986 | 0.943 | 0.943 | 0.953 | 0.947 | 0.946 | 0.920 | 0.920 | 0.920 | 0.923 | 0.918 | 0.926 | 0.926 |
| Cm VQ28 | 1.000 | 1.000 | 0.992 | 0.992 | 0.992 | 0.991 | 0.991 | 0.992 | 0.989 | 0.985 | 0.985 | 0.943 | 0.942 | 0.952 | 0.947 | 0.947 | 0.918 | 0.918 | 0.918 | 0.923 | 0.917 | 0.926 | 0.925 |
| Cm OP3 | 0.999 | 1.000 | 0.991 | 0.991 | 0.992 | 0.991 | 0.991 | 0.991 | 0.989 | 0.984 | 0.984 | 0.942 | 0.942 | 0.953 | 0.946 | 0.946 | 0.918 | 0.918 | 0.918 | 0.923 | 0.917 | 0.926 | 0.926 |
| Cm MAI1001 | 0.991 | 0.991 | 1.000 | 0.997 | 0.995 | 0.990 | 0.991 | 0.990 | 0.988 | 0.984 | 0.984 | 0.944 | 0.943 | 0.952 | 0.947 | 0.947 | 0.917 | 0.918 | 0.918 | 0.923 | 0.918 | 0.926 | 0.925 |
| Cm MAI1050 | 0.991 | 0.990 | 0.996 | 1.000 | 0.996 | 0.991 | 0.991 | 0.990 | 0.988 | 0.985 | 0.984 | 0.942 | 0.942 | 0.952 | 0.946 | 0.946 | 0.918 | 0.918 | 0.918 | 0.923 | 0.918 | 0.926 | 0.926 |
| Cm NT20-18 | 0.992 | 0.992 | 0.995 | 0.996 | 1.000 | 0.991 | 0.992 | 0.991 | 0.989 | 0.985 | 0.985 | 0.943 | 0.943 | 0.953 | 0.947 | 0.947 | 0.918 | 0.918 | 0.918 | 0.924 | 0.918 | 0.927 | 0.926 |
| Cm 0317 | 0.991 | 0.991 | 0.991 | 0.991 | 0.992 | 1.000 | 0.992 | 0.991 | 0.989 | 0.985 | 0.985 | 0.944 | 0.943 | 0.953 | 0.947 | 0.947 | 0.918 | 0.918 | 0.918 | 0.923 | 0.918 | 0.926 | 0.926 |
| Cm NZ2541 | 0.992 | 0.991 | 0.991 | 0.992 | 0.992 | 0.992 | 1.000 | 0.991 | 0.990 | 0.985 | 0.985 | 0.944 | 0.943 | 0.953 | 0.946 | 0.946 | 0.918 | 0.918 | 0.918 | 0.923 | 0.917 | 0.926 | 0.926 |
| Cm CASJ006 | 0.992 | 0.992 | 0.991 | 0.991 | 0.992 | 0.991 | 0.992 | 1.000 | 0.997 | 0.985 | 0.985 | 0.944 | 0.943 | 0.953 | 0.946 | 0.946 | 0.918 | 0.918 | 0.918 | 0.923 | 0.917 | 0.926 | 0.926 |
| Cm CASJ008 | 0.989 | 0.989 | 0.988 | 0.988 | 0.989 | 0.988 | 0.990 | 0.997 | 1.000 | 0.982 | 0.982 | 0.942 | 0.942 | 0.951 | 0.945 | 0.945 | 0.917 | 0.917 | 0.917 | 0.923 | 0.916 | 0.924 | 0.924 |
| Cm Z001 | 0.985 | 0.985 | 0.985 | 0.985 | 0.986 | 0.986 | 0.986 | 0.985 | 0.983 | 1.000 | 1.000 | 0.944 | 0.943 | 0.954 | 0.947 | 0.947 | 0.920 | 0.920 | 0.920 | 0.922 | 0.917 | 0.926 | 0.926 |
| Cm Z002 | 0.985 | 0.985 | 0.985 | 0.985 | 0.986 | 0.985 | 0.986 | 0.985 | 0.983 | 1.000 | 1.000 | 0.944 | 0.943 | 0.953 | 0.947 | 0.947 | 0.920 | 0.920 | 0.920 | 0.922 | 0.917 | 0.926 | 0.926 |
| Ccal CFBP8216 | 0.944 | 0.943 | 0.944 | 0.943 | 0.945 | 0.944 | 0.945 | 0.944 | 0.942 | 0.944 | 0.945 | 1.000 | 0.988 | 0.953 | 0.944 | 0.944 | 0.917 | 0.917 | 0.917 | 0.921 | 0.917 | 0.921 | 0.921 |
| C sp AY1B2 | 0.943 | 0.942 | 0.942 | 0.942 | 0.943 | 0.943 | 0.944 | 0.943 | 0.942 | 0.943 | 0.943 | 0.987 | 1.000 | 0.952 | 0.943 | 0.944 | 0.916 | 0.916 | 0.916 | 0.921 | 0.916 | 0.920 | 0.920 |
| C sp VKM Ac-2542 | 0.952 | 0.952 | 0.952 | 0.952 | 0.953 | 0.953 | 0.953 | 0.953 | 0.951 | 0.953 | 0.953 | 0.953 | 0.952 | 1.000 | 0.954 | 0.954 | 0.920 | 0.920 | 0.920 | 0.924 | 0.920 | 0.926 | 0.926 |
| C sp A6099 | 0.947 | 0.947 | 0.947 | 0.947 | 0.948 | 0.948 | 0.948 | 0.947 | 0.946 | 0.948 | 0.948 | 0.944 | 0.944 | 0.955 | 1.000 | 0.999 | 0.919 | 0.919 | 0.919 | 0.924 | 0.919 | 0.925 | 0.925 |
| C sp LMG 26808 | 0.947 | 0.946 | 0.947 | 0.946 | 0.948 | 0.947 | 0.948 | 0.947 | 0.945 | 0.947 | 0.947 | 0.944 | 0.944 | 0.955 | 0.999 | 1.000 | 0.919 | 0.919 | 0.919 | 0.923 | 0.919 | 0.924 | 0.923 |
| Csep CFIA-Cs3N | 0.918 | 0.918 | 0.918 | 0.918 | 0.919 | 0.918 | 0.919 | 0.918 | 0.917 | 0.920 | 0.920 | 0.917 | 0.917 | 0.920 | 0.919 | 0.919 | 1.000 | 1.000 | 1.000 | 0.912 | 0.911 | 0.918 | 0.917 |
| Csep CFIA-CsR14 | 0.918 | 0.918 | 0.917 | 0.917 | 0.919 | 0.918 | 0.919 | 0.918 | 0.917 | 0.920 | 0.920 | 0.917 | 0.917 | 0.920 | 0.919 | 0.919 | 1.000 | 1.000 | 1.000 | 0.912 | 0.911 | 0.918 | 0.917 |
| Csep ATCC33113 | 0.918 | 0.918 | 0.918 | 0.918 | 0.919 | 0.918 | 0.919 | 0.918 | 0.917 | 0.921 | 0.921 | 0.917 | 0.917 | 0.921 | 0.919 | 0.919 | 1.000 | 1.000 | 1.000 | 0.913 | 0.911 | 0.918 | 0.917 |
| C sp VKM Ac-2872 | 0.923 | 0.923 | 0.923 | 0.923 | 0.925 | 0.923 | 0.925 | 0.923 | 0.923 | 0.923 | 0.923 | 0.921 | 0.921 | 0.925 | 0.923 | 0.923 | 0.912 | 0.912 | 0.912 | 1.000 | 0.933 | 0.924 | 0.924 |
| C sp VKM Ac-2873 | 0.918 | 0.918 | 0.918 | 0.918 | 0.920 | 0.919 | 0.920 | 0.918 | 0.918 | 0.918 | 0.918 | 0.917 | 0.918 | 0.920 | 0.919 | 0.920 | 0.910 | 0.910 | 0.910 | 0.934 | 1.000 | 0.920 | 0.920 |
| Cins CFBP2404 | 0.927 | 0.927 | 0.927 | 0.926 | 0.928 | 0.927 | 0.928 | 0.927 | 0.926 | 0.927 | 0.927 | 0.921 | 0.921 | 0.927 | 0.924 | 0.924 | 0.917 | 0.917 | 0.917 | 0.924 | 0.920 | 1.000 | 1.000 |
| Cins LMG 3663 | 0.926 | 0.926 | 0.926 | 0.926 | 0.928 | 0.926 | 0.927 | 0.926 | 0.925 | 0.926 | 0.927 | 0.921 | 0.921 | 0.927 | 0.924 | 0.924 | 0.917 | 0.917 | 0.917 | 0.925 | 0.920 | 1.000 | 1.000 |
| Cins R1-1 | 0.926 | 0.926 | 0.926 | 0.926 | 0.928 | 0.926 | 0.927 | 0.926 | 0.925 | 0.926 | 0.926 | 0.921 | 0.921 | 0.926 | 0.924 | 0.924 | 0.917 | 0.917 | 0.917 | 0.924 | 0.920 | 0.998 | 0.998 |
| Cins R1-3 | 0.926 | 0.926 | 0.926 | 0.926 | 0.929 | 0.926 | 0.928 | 0.926 | 0.926 | 0.927 | 0.927 | 0.921 | 0.921 | 0.927 | 0.924 | 0.924 | 0.918 | 0.918 | 0.918 | 0.925 | 0.920 | 0.999 | 0.999 |
| C sp CFBP7494 | 0.929 | 0.929 | 0.928 | 0.928 | 0.930 | 0.929 | 0.930 | 0.928 | 0.928 | 0.929 | 0.929 | 0.924 | 0.925 | 0.929 | 0.926 | 0.926 | 0.919 | 0.919 | 0.919 | 0.927 | 0.923 | 0.957 | 0.957 |
| Cneb 61-1 | 0.925 | 0.925 | 0.925 | 0.925 | 0.927 | 0.925 | 0.927 | 0.925 | 0.925 | 0.926 | 0.926 | 0.921 | 0.922 | 0.925 | 0.923 | 0.923 | 0.918 | 0.918 | 0.918 | 0.924 | 0.919 | 0.950 | 0.950 |
| Cneb 7580 | 0.925 | 0.925 | 0.926 | 0.925 | 0.927 | 0.926 | 0.927 | 0.925 | 0.925 | 0.926 | 0.926 | 0.921 | 0.922 | 0.925 | 0.923 | 0.923 | 0.918 | 0.918 | 0.918 | 0.924 | 0.919 | 0.950 | 0.950 |
| Cneb DOAB 395 | 0.925 | 0.925 | 0.926 | 0.925 | 0.927 | 0.926 | 0.926 | 0.925 | 0.924 | 0.926 | 0.926 | 0.921 | 0.921 | 0.925 | 0.923 | 0.923 | 0.919 | 0.919 | 0.919 | 0.924 | 0.920 | 0.950 | 0.950 |
| Cneb NCPPB 2581 | 0.925 | 0.925 | 0.925 | 0.925 | 0.927 | 0.925 | 0.926 | 0.925 | 0.924 | 0.926 | 0.926 | 0.921 | 0.921 | 0.925 | 0.923 | 0.923 | 0.918 | 0.918 | 0.918 | 0.923 | 0.920 | 0.950 | 0.950 |
| C sp PvP098 | 0.930 | 0.930 | 0.930 | 0.930 | 0.932 | 0.930 | 0.931 | 0.930 | 0.929 | 0.931 | 0.931 | 0.925 | 0.925 | 0.931 | 0.928 | 0.928 | 0.922 | 0.923 | 0.923 | 0.926 | 0.922 | 0.937 | 0.937 |
| Cchil CFBP8217 | 0.925 | 0.924 | 0.925 | 0.924 | 0.926 | 0.925 | 0.926 | 0.925 | 0.925 | 0.926 | 0.926 | 0.920 | 0.921 | 0.923 | 0.920 | 0.920 | 0.917 | 0.917 | 0.917 | 0.921 | 0.920 | 0.928 | 0.928 |
| Cphas LPPA 982 | 0.924 | 0.924 | 0.924 | 0.924 | 0.926 | 0.925 | 0.926 | 0.925 | 0.924 | 0.925 | 0.925 | 0.919 | 0.920 | 0.922 | 0.920 | 0.920 | 0.917 | 0.917 | 0.917 | 0.920 | 0.919 | 0.927 | 0.927 |
| C sp PvP097 | 0.925 | 0.925 | 0.925 | 0.925 | 0.927 | 0.925 | 0.926 | 0.925 | 0.924 | 0.926 | 0.926 | 0.920 | 0.920 | 0.923 | 0.921 | 0.921 | 0.917 | 0.918 | 0.918 | 0.922 | 0.921 | 0.929 | 0.928 |
| Cphas VKM Ac-2886 | 0.923 | 0.923 | 0.923 | 0.923 | 0.925 | 0.923 | 0.925 | 0.923 | 0.923 | 0.924 | 0.924 | 0.918 | 0.919 | 0.921 | 0.919 | 0.920 | 0.914 | 0.915 | 0.915 | 0.919 | 0.918 | 0.926 | 0.926 |
| C sp PvP036 | 0.922 | 0.922 | 0.922 | 0.922 | 0.924 | 0.923 | 0.924 | 0.922 | 0.922 | 0.923 | 0.923 | 0.918 | 0.918 | 0.921 | 0.919 | 0.919 | 0.916 | 0.916 | 0.916 | 0.919 | 0.918 | 0.926 | 0.926 |
| C sp CFBP8019 | 0.911 | 0.911 | 0.911 | 0.911 | 0.913 | 0.911 | 0.913 | 0.911 | 0.911 | 0.912 | 0.912 | 0.909 | 0.909 | 0.911 | 0.910 | 0.910 | 0.909 | 0.909 | 0.909 | 0.908 | 0.907 | 0.912 | 0.912 |
| C sp CFBP3399 | 0.892 | 0.892 | 0.892 | 0.893 | 0.894 | 0.892 | 0.894 | 0.892 | 0.893 | 0.893 | 0.893 | 0.891 | 0.891 | 0.891 | 0.891 | 0.891 | 0.887 | 0.887 | 0.887 | 0.892 | 0.890 | 0.894 | 0.894 |
| Ctes ATCC 33566 | 0.892 | 0.892 | 0.892 | 0.893 | 0.895 | 0.892 | 0.894 | 0.892 | 0.892 | 0.893 | 0.893 | 0.890 | 0.890 | 0.891 | 0.890 | 0.891 | 0.887 | 0.887 | 0.887 | 0.891 | 0.890 | 0.894 | 0.893 |
| C sp CFBP8017 | 0.889 | 0.889 | 0.890 | 0.890 | 0.892 | 0.890 | 0.892 | 0.890 | 0.890 | 0.890 | 0.890 | 0.889 | 0.889 | 0.889 | 0.888 | 0.888 | 0.886 | 0.885 | 0.886 | 0.890 | 0.888 | 0.890 | 0.890 |
| Ctes DOAB 609 | 0.891 | 0.891 | 0.891 | 0.891 | 0.893 | 0.891 | 0.893 | 0.891 | 0.892 | 0.891 | 0.891 | 0.890 | 0.890 | 0.890 | 0.890 | 0.891 | 0.887 | 0.887 | 0.887 | 0.890 | 0.889 | 0.891 | 0.891 |
| Czhang DM1 | 0.894 | 0.894 | 0.894 | 0.894 | 0.896 | 0.894 | 0.896 | 0.894 | 0.894 | 0.894 | 0.894 | 0.893 | 0.893 | 0.894 | 0.893 | 0.894 | 0.891 | 0.891 | 0.891 | 0.892 | 0.892 | 0.896 | 0.895 |
| Ccap PF008 | 0.902 | 0.902 | 0.901 | 0.901 | 0.904 | 0.902 | 0.903 | 0.902 | 0.902 | 0.902 | 0.902 | 0.899 | 0.899 | 0.901 | 0.900 | 0.900 | 0.898 | 0.898 | 0.898 | 0.899 | 0.898 | 0.903 | 0.902 |
| Ccap RA1B | 0.903 | 0.903 | 0.903 | 0.903 | 0.904 | 0.903 | 0.904 | 0.903 | 0.903 | 0.903 | 0.903 | 0.900 | 0.900 | 0.902 | 0.901 | 0.902 | 0.898 | 0.899 | 0.899 | 0.900 | 0.898 | 0.904 | 0.904 |
| Ccap 1207 | 0.902 | 0.902 | 0.902 | 0.902 | 0.904 | 0.902 | 0.904 | 0.903 | 0.903 | 0.903 | 0.903 | 0.899 | 0.899 | 0.902 | 0.901 | 0.901 | 0.898 | 0.899 | 0.899 | 0.899 | 0.898 | 0.903 | 0.903 |
| C sp AY1A6 | 0.880 | 0.880 | 0.880 | 0.880 | 0.882 | 0.880 | 0.882 | 0.880 | 0.881 | 0.880 | 0.881 | 0.881 | 0.882 | 0.881 | 0.879 | 0.880 | 0.878 | 0.878 | 0.878 | 0.880 | 0.879 | 0.882 | 0.881 |
| C sp CASJ009 | 0.880 | 0.880 | 0.880 | 0.880 | 0.882 | 0.880 | 0.882 | 0.880 | 0.881 | 0.880 | 0.880 | 0.879 | 0.880 | 0.880 | 0.880 | 0.880 | 0.879 | 0.878 | 0.878 | 0.879 | 0.878 | 0.881 | 0.881 |
| C sp AY1B3 | 0.880 | 0.880 | 0.880 | 0.880 | 0.882 | 0.880 | 0.882 | 0.880 | 0.881 | 0.880 | 0.880 | 0.880 | 0.882 | 0.880 | 0.879 | 0.880 | 0.878 | 0.878 | 0.878 | 0.880 | 0.879 | 0.881 | 0.880 |

| | Cins R1-1 | Cins R1-3 | C sp CFBP7494 | Cneb 61-1 | Cneb 7580 | Cneb DOAB 395 | Cneb NCPPB 2581 | C sp PvP098 | Cchil CFBP8217 | Cphas LPPA 982 | C sp PvP097 | Cphas VKM Ac-2886 | C sp PvP036 | C sp CFBP8019 | C sp CFBP3399 | Ctes ATCC 33566 | C sp CFBP8017 | Ctes DOAB 609 | Czhang DM1 | Ccap PF008 | Ccap RA1B | Ccap 1207 | C sp AY1A6 | C sp CASJ009 | C sp AY1B3 |
|---|---|---|---|---|---|---|---|---|---|---|---|---|---|---|---|---|---|---|---|---|---|---|---|---|---|
| Cm CFBP5842 | 0.924 | 0.925 | 0.928 | 0.923 | 0.923 | 0.924 | 0.923 | 0.929 | 0.924 | 0.924 | 0.924 | 0.923 | 0.921 | 0.911 | 0.891 | 0.891 | 0.889 | 0.889 | 0.894 | 0.900 | 0.902 | 0.900 | 0.881 | 0.879 | 0.880 |
| Cm CMM84 | 0.925 | 0.925 | 0.929 | 0.924 | 0.924 | 0.924 | 0.924 | 0.929 | 0.924 | 0.924 | 0.924 | 0.924 | 0.922 | 0.911 | 0.892 | 0.892 | 0.890 | 0.890 | 0.894 | 0.900 | 0.903 | 0.900 | 0.882 | 0.880 | 0.881 |
| Cm ATCC 14456 | 0.925 | 0.925 | 0.928 | 0.923 | 0.923 | 0.923 | 0.923 | 0.929 | 0.924 | 0.924 | 0.924 | 0.923 | 0.922 | 0.911 | 0.891 | 0.891 | 0.889 | 0.889 | 0.894 | 0.901 | 0.903 | 0.901 | 0.881 | 0.879 | 0.880 |
| Cm CFBP7568 | 0.925 | 0.925 | 0.929 | 0.924 | 0.924 | 0.924 | 0.924 | 0.929 | 0.924 | 0.924 | 0.924 | 0.923 | 0.922 | 0.910 | 0.891 | 0.891 | 0.890 | 0.889 | 0.893 | 0.901 | 0.902 | 0.901 | 0.881 | 0.880 | 0.880 |
| Cm MX14-E106-3 | 0.924 | 0.924 | 0.927 | 0.923 | 0.923 | 0.923 | 0.923 | 0.928 | 0.923 | 0.923 | 0.923 | 0.922 | 0.920 | 0.910 | 0.891 | 0.891 | 0.889 | 0.889 | 0.893 | 0.900 | 0.902 | 0.900 | 0.881 | 0.879 | 0.879 |
| Cm MX16-H8B | 0.924 | 0.924 | 0.928 | 0.923 | 0.923 | 0.923 | 0.923 | 0.929 | 0.924 | 0.924 | 0.924 | 0.923 | 0.921 | 0.910 | 0.891 | 0.891 | 0.889 | 0.889 | 0.894 | 0.899 | 0.902 | 0.899 | 0.881 | 0.879 | 0.880 |
| Cm LMG7333 | 0.925 | 0.926 | 0.929 | 0.924 | 0.924 | 0.924 | 0.924 | 0.929 | 0.924 | 0.924 | 0.924 | 0.923 | 0.922 | 0.911 | 0.893 | 0.893 | 0.890 | 0.890 | 0.895 | 0.901 | 0.903 | 0.901 | 0.881 | 0.880 | 0.880 |
| Cm VKM Ac-1790 | 0.926 | 0.926 | 0.928 | 0.924 | 0.924 | 0.924 | 0.924 | 0.929 | 0.925 | 0.925 | 0.924 | 0.924 | 0.922 | 0.912 | 0.892 | 0.892 | 0.890 | 0.890 | 0.894 | 0.901 | 0.903 | 0.901 | 0.881 | 0.879 | 0.880 |
| Cm CFBP7314 | 0.925 | 0.925 | 0.929 | 0.924 | 0.924 | 0.924 | 0.924 | 0.929 | 0.924 | 0.924 | 0.924 | 0.923 | 0.922 | 0.911 | 0.891 | 0.891 | 0.889 | 0.889 | 0.893 | 0.901 | 0.903 | 0.901 | 0.881 | 0.880 | 0.880 |
| Cm 1217 | 0.926 | 0.926 | 0.930 | 0.925 | 0.925 | 0.925 | 0.925 | 0.930 | 0.926 | 0.926 | 0.925 | 0.924 | 0.923 | 0.913 | 0.894 | 0.894 | 0.891 | 0.891 | 0.896 | 0.902 | 0.903 | 0.902 | 0.882 | 0.881 | 0.881 |
| Cm MX15-L3A | 0.925 | 0.925 | 0.928 | 0.924 | 0.924 | 0.924 | 0.924 | 0.929 | 0.924 | 0.924 | 0.924 | 0.922 | 0.921 | 0.911 | 0.892 | 0.892 | 0.890 | 0.889 | 0.894 | 0.900 | 0.903 | 0.900 | 0.882 | 0.880 | 0.880 |
| Cm MX16-I12A | 0.925 | 0.926 | 0.928 | 0.923 | 0.923 | 0.923 | 0.923 | 0.929 | 0.924 | 0.924 | 0.924 | 0.923 | 0.921 | 0.911 | 0.891 | 0.891 | 0.890 | 0.889 | 0.894 | 0.900 | 0.902 | 0.900 | 0.881 | 0.880 | 0.880 |
| Cm CFBP7158 | 0.925 | 0.925 | 0.928 | 0.923 | 0.923 | 0.924 | 0.923 | 0.929 | 0.924 | 0.924 | 0.924 | 0.923 | 0.921 | 0.911 | 0.891 | 0.891 | 0.890 | 0.889 | 0.894 | 0.900 | 0.902 | 0.901 | 0.881 | 0.879 | 0.879 |
| Cm NCPPB 382 | 0.925 | 0.926 | 0.929 | 0.924 | 0.924 | 0.924 | 0.924 | 0.929 | 0.924 | 0.924 | 0.924 | 0.923 | 0.922 | 0.911 | 0.892 | 0.892 | 0.890 | 0.890 | 0.894 | 0.901 | 0.903 | 0.901 | 0.881 | 0.879 | 0.880 |
| Cm MX13-E96-2 | 0.925 | 0.925 | 0.928 | 0.924 | 0.924 | 0.924 | 0.924 | 0.929 | 0.924 | 0.924 | 0.924 | 0.923 | 0.922 | 0.911 | 0.892 | 0.891 | 0.890 | 0.889 | 0.894 | 0.900 | 0.902 | 0.900 | 0.881 | 0.880 | 0.880 |
| Cm MAI1009 | 0.925 | 0.926 | 0.929 | 0.924 | 0.924 | 0.924 | 0.924 | 0.930 | 0.925 | 0.925 | 0.925 | 0.923 | 0.922 | 0.912 | 0.893 | 0.892 | 0.890 | 0.890 | 0.894 | 0.901 | 0.903 | 0.901 | 0.882 | 0.880 | 0.881 |
| Cm MSF322 | 0.925 | 0.926 | 0.929 | 0.924 | 0.924 | 0.924 | 0.924 | 0.930 | 0.925 | 0.925 | 0.925 | 0.924 | 0.922 | 0.912 | 0.892 | 0.892 | 0.890 | 0.890 | 0.895 | 0.900 | 0.903 | 0.901 | 0.881 | 0.880 | 0.881 |
| Cm CFBP7315 | 0.926 | 0.926 | 0.929 | 0.924 | 0.924 | 0.924 | 0.924 | 0.929 | 0.924 | 0.924 | 0.924 | 0.923 | 0.922 | 0.911 | 0.892 | 0.892 | 0.890 | 0.890 | 0.894 | 0.901 | 0.903 | 0.901 | 0.881 | 0.880 | 0.881 |
| Cm VL527 | 0.925 | 0.925 | 0.928 | 0.924 | 0.924 | 0.924 | 0.924 | 0.929 | 0.924 | 0.924 | 0.924 | 0.923 | 0.922 | 0.911 | 0.891 | 0.891 | 0.889 | 0.890 | 0.894 | 0.900 | 0.903 | 0.901 | 0.881 | 0.880 | 0.881 |
| Cm MX15-115 | 0.924 | 0.925 | 0.928 | 0.924 | 0.924 | 0.924 | 0.924 | 0.929 | 0.924 | 0.924 | 0.924 | 0.923 | 0.922 | 0.911 | 0.892 | 0.892 | 0.890 | 0.890 | 0.895 | 0.901 | 0.903 | 0.901 | 0.881 | 0.879 | 0.880 |
| Cm VQ143 | 0.926 | 0.926 | 0.929 | 0.924 | 0.924 | 0.924 | 0.924 | 0.930 | 0.925 | 0.925 | 0.925 | 0.924 | 0.923 | 0.912 | 0.892 | 0.892 | 0.890 | 0.890 | 0.894 | 0.901 | 0.903 | 0.901 | 0.882 | 0.881 | 0.882 |
| Cm VQ28 | 0.925 | 0.925 | 0.929 | 0.923 | 0.923 | 0.924 | 0.923 | 0.929 | 0.924 | 0.924 | 0.924 | 0.923 | 0.922 | 0.911 | 0.891 | 0.892 | 0.890 | 0.889 | 0.894 | 0.901 | 0.903 | 0.901 | 0.881 | 0.880 | 0.880 |
| Cm OP3 | 0.926 | 0.926 | 0.929 | 0.924 | 0.924 | 0.924 | 0.924 | 0.930 | 0.924 | 0.924 | 0.925 | 0.923 | 0.922 | 0.911 | 0.892 | 0.892 | 0.890 | 0.890 | 0.894 | 0.901 | 0.903 | 0.900 | 0.881 | 0.880 | 0.881 |
| Cm MAI1001 | 0.925 | 0.925 | 0.928 | 0.924 | 0.924 | 0.924 | 0.924 | 0.930 | 0.925 | 0.925 | 0.925 | 0.924 | 0.922 | 0.911 | 0.892 | 0.892 | 0.890 | 0.890 | 0.894 | 0.900 | 0.902 | 0.901 | 0.881 | 0.880 | 0.880 |
| Cm MAI1050 | 0.926 | 0.926 | 0.928 | 0.924 | 0.924 | 0.924 | 0.924 | 0.929 | 0.924 | 0.924 | 0.924 | 0.923 | 0.922 | 0.911 | 0.892 | 0.893 | 0.890 | 0.890 | 0.894 | 0.900 | 0.902 | 0.900 | 0.881 | 0.880 | 0.880 |
| Cm NT20-18 | 0.926 | 0.926 | 0.929 | 0.925 | 0.925 | 0.925 | 0.925 | 0.930 | 0.924 | 0.924 | 0.925 | 0.924 | 0.922 | 0.912 | 0.893 | 0.893 | 0.891 | 0.891 | 0.895 | 0.902 | 0.903 | 0.901 | 0.882 | 0.881 | 0.881 |
| Cm 0317 | 0.926 | 0.926 | 0.929 | 0.924 | 0.924 | 0.924 | 0.924 | 0.930 | 0.925 | 0.925 | 0.924 | 0.923 | 0.921 | 0.911 | 0.892 | 0.891 | 0.890 | 0.889 | 0.894 | 0.900 | 0.902 | 0.901 | 0.881 | 0.879 | 0.880 |
| Cm NZ2541 | 0.925 | 0.925 | 0.928 | 0.924 | 0.924 | 0.924 | 0.924 | 0.929 | 0.924 | 0.924 | 0.924 | 0.923 | 0.921 | 0.911 | 0.892 | 0.892 | 0.889 | 0.890 | 0.894 | 0.900 | 0.902 | 0.900 | 0.881 | 0.879 | 0.880 |
| Cm CASJ006 | 0.925 | 0.925 | 0.928 | 0.923 | 0.923 | 0.923 | 0.923 | 0.929 | 0.924 | 0.924 | 0.924 | 0.923 | 0.921 | 0.911 | 0.891 | 0.891 | 0.889 | 0.890 | 0.893 | 0.901 | 0.903 | 0.901 | 0.881 | 0.880 | 0.881 |
| Cm CASJ008 | 0.924 | 0.924 | 0.927 | 0.923 | 0.923 | 0.923 | 0.923 | 0.928 | 0.924 | 0.924 | 0.923 | 0.922 | 0.921 | 0.910 | 0.892 | 0.892 | 0.890 | 0.890 | 0.895 | 0.901 | 0.903 | 0.901 | 0.881 | 0.880 | 0.880 |
| Cm Z001 | 0.926 | 0.926 | 0.929 | 0.924 | 0.924 | 0.925 | 0.924 | 0.930 | 0.925 | 0.925 | 0.925 | 0.924 | 0.922 | 0.911 | 0.892 | 0.893 | 0.890 | 0.890 | 0.894 | 0.901 | 0.903 | 0.901 | 0.881 | 0.880 | 0.881 |
| Cm Z002 | 0.926 | 0.926 | 0.929 | 0.924 | 0.924 | 0.924 | 0.924 | 0.930 | 0.924 | 0.925 | 0.924 | 0.924 | 0.922 | 0.911 | 0.892 | 0.892 | 0.890 | 0.889 | 0.894 | 0.901 | 0.902 | 0.901 | 0.881 | 0.879 | 0.880 |
| Ccal CFBP8216 | 0.921 | 0.921 | 0.924 | 0.921 | 0.921 | 0.921 | 0.921 | 0.925 | 0.920 | 0.920 | 0.919 | 0.919 | 0.917 | 0.910 | 0.891 | 0.891 | 0.889 | 0.890 | 0.894 | 0.899 | 0.902 | 0.900 | 0.883 | 0.880 | 0.883 |
| C sp AY1B2 | 0.920 | 0.920 | 0.923 | 0.920 | 0.920 | 0.920 | 0.920 | 0.924 | 0.919 | 0.919 | 0.918 | 0.918 | 0.916 | 0.909 | 0.891 | 0.890 | 0.889 | 0.889 | 0.893 | 0.898 | 0.901 | 0.898 | 0.882 | 0.879 | 0.882 |
| C sp VKM Ac-2542 | 0.925 | 0.925 | 0.928 | 0.924 | 0.924 | 0.924 | 0.924 | 0.931 | 0.922 | 0.922 | 0.922 | 0.921 | 0.920 | 0.910 | 0.891 | 0.891 | 0.889 | 0.890 | 0.894 | 0.900 | 0.902 | 0.900 | 0.882 | 0.880 | 0.881 |
| C sp A6099 | 0.924 | 0.925 | 0.927 | 0.923 | 0.923 | 0.923 | 0.923 | 0.928 | 0.921 | 0.921 | 0.921 | 0.920 | 0.919 | 0.911 | 0.891 | 0.891 | 0.889 | 0.889 | 0.894 | 0.900 | 0.902 | 0.900 | 0.882 | 0.880 | 0.881 |
| C sp LMG 26808 | 0.923 | 0.923 | 0.926 | 0.922 | 0.922 | 0.922 | 0.922 | 0.928 | 0.920 | 0.920 | 0.920 | 0.919 | 0.918 | 0.910 | 0.891 | 0.891 | 0.888 | 0.889 | 0.894 | 0.900 | 0.901 | 0.900 | 0.881 | 0.879 | 0.881 |
| Csep CFIA-Cs3N | 0.917 | 0.917 | 0.920 | 0.918 | 0.918 | 0.918 | 0.918 | 0.923 | 0.917 | 0.917 | 0.917 | 0.916 | 0.916 | 0.909 | 0.887 | 0.888 | 0.887 | 0.886 | 0.892 | 0.899 | 0.900 | 0.898 | 0.880 | 0.879 | 0.879 |
| Csep CFIA-CsR14 | 0.916 | 0.917 | 0.920 | 0.917 | 0.917 | 0.917 | 0.917 | 0.923 | 0.917 | 0.917 | 0.917 | 0.915 | 0.916 | 0.909 | 0.887 | 0.887 | 0.886 | 0.885 | 0.891 | 0.898 | 0.900 | 0.898 | 0.879 | 0.878 | 0.878 |
| Csep ATCC33113 | 0.918 | 0.918 | 0.921 | 0.918 | 0.918 | 0.918 | 0.918 | 0.923 | 0.917 | 0.917 | 0.917 | 0.916 | 0.916 | 0.909 | 0.888 | 0.888 | 0.887 | 0.886 | 0.892 | 0.899 | 0.900 | 0.899 | 0.880 | 0.879 | 0.880 |
| C sp VKM Ac-2872 | 0.924 | 0.924 | 0.928 | 0.923 | 0.923 | 0.923 | 0.923 | 0.926 | 0.921 | 0.921 | 0.921 | 0.920 | 0.919 | 0.908 | 0.891 | 0.892 | 0.890 | 0.888 | 0.893 | 0.898 | 0.900 | 0.897 | 0.880 | 0.879 | 0.880 |
| C sp VKM Ac-2873 | 0.920 | 0.920 | 0.923 | 0.919 | 0.919 | 0.920 | 0.919 | 0.922 | 0.920 | 0.920 | 0.920 | 0.919 | 0.918 | 0.908 | 0.890 | 0.889 | 0.888 | 0.888 | 0.892 | 0.898 | 0.900 | 0.898 | 0.880 | 0.879 | 0.879 |
| Cins CFBP2404 | 0.998 | 0.998 | 0.957 | 0.950 | 0.950 | 0.950 | 0.950 | 0.937 | 0.928 | 0.928 | 0.928 | 0.927 | 0.926 | 0.913 | 0.893 | 0.893 | 0.892 | 0.891 | 0.896 | 0.903 | 0.905 | 0.903 | 0.882 | 0.881 | 0.882 |
| Cins LMG 3663 | 0.998 | 0.998 | 0.957 | 0.950 | 0.950 | 0.950 | 0.950 | 0.937 | 0.928 | 0.928 | 0.928 | 0.926 | 0.926 | 0.912 | 0.893 | 0.893 | 0.891 | 0.891 | 0.896 | 0.903 | 0.905 | 0.903 | 0.883 | 0.881 | 0.882 |
| Cins R1-1 | 1.000 | 0.999 | 0.957 | 0.949 | 0.949 | 0.949 | 0.949 | 0.936 | 0.927 | 0.927 | 0.927 | 0.926 | 0.925 | 0.912 | 0.893 | 0.893 | 0.891 | 0.891 | 0.896 | 0.902 | 0.904 | 0.902 | 0.883 | 0.881 | 0.882 |
| Cins R1-3 | 1.000 | 1.000 | 0.957 | 0.949 | 0.949 | 0.950 | 0.949 | 0.937 | 0.928 | 0.928 | 0.928 | 0.927 | 0.926 | 0.913 | 0.893 | 0.893 | 0.892 | 0.892 | 0.896 | 0.903 | 0.905 | 0.903 | 0.882 | 0.881 | 0.882 |
| C sp CFBP7494 | 0.957 | 0.957 | 1.000 | 0.950 | 0.950 | 0.950 | 0.950 | 0.941 | 0.930 | 0.930 | 0.930 | 0.929 | 0.928 | 0.914 | 0.896 | 0.896 | 0.893 | 0.892 | 0.896 | 0.904 | 0.906 | 0.904 | 0.883 | 0.881 | 0.882 |
| Cneb 61-1 | 0.950 | 0.950 | 0.950 | 1.000 | 1.000 | 1.000 | 1.000 | 0.935 | 0.930 | 0.930 | 0.929 | 0.929 | 0.927 | 0.913 | 0.894 | 0.893 | 0.890 | 0.891 | 0.896 | 0.904 | 0.906 | 0.905 | 0.883 | 0.882 | 0.882 |
| Cneb 7580 | 0.950 | 0.950 | 0.950 | 1.000 | 1.000 | 1.000 | 1.000 | 0.935 | 0.930 | 0.930 | 0.929 | 0.929 | 0.927 | 0.913 | 0.894 | 0.893 | 0.890 | 0.891 | 0.896 | 0.904 | 0.906 | 0.905 | 0.883 | 0.882 | 0.882 |
| Cneb DOAB 395 | 0.950 | 0.950 | 0.951 | 1.000 | 1.000 | 1.000 | 1.000 | 0.935 | 0.930 | 0.930 | 0.929 | 0.929 | 0.927 | 0.913 | 0.894 | 0.894 | 0.891 | 0.892 | 0.897 | 0.905 | 0.907 | 0.905 | 0.883 | 0.882 | 0.883 |
| Cneb NCPPB 2581 | 0.950 | 0.950 | 0.950 | 1.000 | 1.000 | 1.000 | 1.000 | 0.935 | 0.929 | 0.929 | 0.929 | 0.929 | 0.927 | 0.913 | 0.894 | 0.893 | 0.890 | 0.891 | 0.897 | 0.905 | 0.906 | 0.905 | 0.882 | 0.881 | 0.882 |
| C sp PvP098 | 0.937 | 0.936 | 0.941 | 0.934 | 0.934 | 0.934 | 0.934 | 1.000 | 0.931 | 0.931 | 0.931 | 0.930 | 0.929 | 0.914 | 0.894 | 0.893 | 0.891 | 0.891 | 0.896 | 0.904 | 0.906 | 0.904 | 0.883 | 0.882 | 0.882 |
| Cchil CFBP8217 | 0.928 | 0.928 | 0.930 | 0.930 | 0.930 | 0.930 | 0.930 | 0.931 | 1.000 | 1.000 | 0.983 | 0.973 | 0.953 | 0.914 | 0.894 | 0.893 | 0.892 | 0.893 | 0.897 | 0.906 | 0.909 | 0.907 | 0.884 | 0.882 | 0.884 |
| Cphas LPPA 982 | 0.927 | 0.927 | 0.930 | 0.928 | 0.928 | 0.929 | 0.928 | 0.931 | 1.000 | 1.000 | 0.983 | 0.972 | 0.954 | 0.914 | 0.894 | 0.893 | 0.892 | 0.893 | 0.897 | 0.906 | 0.908 | 0.906 | 0.885 | 0.883 | 0.884 |
| C sp PvP097 | 0.928 | 0.928 | 0.931 | 0.929 | 0.929 | 0.929 | 0.929 | 0.932 | 0.984 | 0.984 | 1.000 | 0.973 | 0.954 | 0.915 | 0.895 | 0.894 | 0.892 | 0.893 | 0.898 | 0.907 | 0.909 | 0.907 | 0.885 | 0.884 | 0.885 |
| Cphas VKM Ac-2886 | 0.926 | 0.926 | 0.928 | 0.928 | 0.928 | 0.928 | 0.928 | 0.929 | 0.972 | 0.972 | 0.972 | 1.000 | 0.953 | 0.914 | 0.893 | 0.892 | 0.891 | 0.891 | 0.896 | 0.905 | 0.907 | 0.905 | 0.883 | 0.883 | 0.884 |
| C sp PvP036 | 0.926 | 0.926 | 0.928 | 0.926 | 0.926 | 0.927 | 0.926 | 0.929 | 0.954 | 0.954 | 0.954 | 0.954 | 1.000 | 0.914 | 0.893 | 0.893 | 0.890 | 0.892 | 0.897 | 0.906 | 0.907 | 0.906 | 0.883 | 0.883 | 0.884 |
| C sp CFBP8019 | 0.912 | 0.912 | 0.914 | 0.911 | 0.911 | 0.911 | 0.911 | 0.914 | 0.914 | 0.914 | 0.914 | 0.914 | 0.913 | 1.000 | 0.891 | 0.890 | 0.889 | 0.890 | 0.896 | 0.902 | 0.903 | 0.901 | 0.881 | 0.880 | 0.881 |
| C sp CFBP3399 | 0.893 | 0.893 | 0.895 | 0.893 | 0.893 | 0.894 | 0.893 | 0.894 | 0.895 | 0.895 | 0.894 | 0.894 | 0.893 | 0.890 | 1.000 | 0.983 | 0.946 | 0.926 | 0.895 | 0.896 | 0.898 | 0.895 | 0.877 | 0.876 | 0.877 |
| Ctes ATCC 33566 | 0.893 | 0.893 | 0.895 | 0.893 | 0.893 | 0.893 | 0.893 | 0.894 | 0.894 | 0.894 | 0.894 | 0.892 | 0.893 | 0.890 | 0.984 | 1.000 | 0.945 | 0.926 | 0.894 | 0.895 | 0.897 | 0.895 | 0.876 | 0.875 | 0.876 |
| C sp CFBP8017 | 0.890 | 0.890 | 0.892 | 0.889 | 0.889 | 0.889 | 0.889 | 0.891 | 0.892 | 0.892 | 0.891 | 0.891 | 0.890 | 0.888 | 0.946 | 0.945 | 1.000 | 0.921 | 0.892 | 0.892 | 0.894 | 0.893 | 0.876 | 0.874 | 0.876 |
| Ctes DOAB 609 | 0.891 | 0.891 | 0.893 | 0.892 | 0.892 | 0.892 | 0.892 | 0.892 | 0.894 | 0.894 | 0.894 | 0.893 | 0.893 | 0.891 | 0.927 | 0.927 | 0.922 | 1.000 | 0.894 | 0.896 | 0.898 | 0.896 | 0.876 | 0.874 | 0.876 |
| Czhang DM1 | 0.895 | 0.895 | 0.896 | 0.896 | 0.896 | 0.896 | 0.896 | 0.895 | 0.897 | 0.897 | 0.897 | 0.897 | 0.897 | 0.896 | 0.895 | 0.894 | 0.893 | 0.894 | 1.000 | 0.904 | 0.905 | 0.904 | 0.880 | 0.879 | 0.880 |
| Ccap PF008 | 0.902 | 0.903 | 0.904 | 0.904 | 0.904 | 0.904 | 0.904 | 0.905 | 0.907 | 0.907 | 0.907 | 0.907 | 0.906 | 0.903 | 0.896 | 0.896 | 0.894 | 0.895 | 0.904 | 1.000 | 0.990 | 0.988 | 0.881 | 0.879 | 0.880 |
| Ccap RA1B | 0.904 | 0.904 | 0.906 | 0.904 | 0.904 | 0.904 | 0.905 | 0.905 | 0.908 | 0.908 | 0.908 | 0.907 | 0.907 | 0.903 | 0.897 | 0.897 | 0.894 | 0.896 | 0.905 | 0.989 | 1.000 | 0.987 | 0.882 | 0.881 | 0.882 |
| Ccap 1207 | 0.903 | 0.903 | 0.906 | 0.905 | 0.905 | 0.905 | 0.905 | 0.905 | 0.907 | 0.907 | 0.908 | 0.907 | 0.906 | 0.902 | 0.896 | 0.896 | 0.895 | 0.896 | 0.904 | 0.988 | 0.988 | 1.000 | 0.882 | 0.881 | 0.882 |
| C sp AY1A6 | 0.881 | 0.881 | 0.882 | 0.881 | 0.881 | 0.882 | 0.881 | 0.882 | 0.884 | 0.884 | 0.883 | 0.882 | 0.882 | 0.880 | 0.876 | 0.875 | 0.875 | 0.875 | 0.879 | 0.880 | 0.882 | 0.880 | 1.000 | 0.982 | 0.983 |
| C sp CASJ009 | 0.881 | 0.881 | 0.882 | 0.881 | 0.881 | 0.881 | 0.881 | 0.881 | 0.883 | 0.883 | 0.883 | 0.882 | 0.882 | 0.880 | 0.877 | 0.876 | 0.875 | 0.875 | 0.880 | 0.880 | 0.882 | 0.880 | 0.983 | 1.000 | 0.983 |
| C sp AY1B3 | 0.880 | 0.880 | 0.881 | 0.881 | 0.881 | 0.881 | 0.881 | 0.881 | 0.883 | 0.883 | 0.884 | 0.884 | 0.883 | 0.881 | 0.876 | 0.875 | 0.875 | 0.875 | 0.879 | 0.880 | 0.882 | 0.880 | 0.984 | 0.982 | 1.000 |

**Supplementary table S3**: *Clavibacter* genus genome database information. *Cm, C. michiganensis; C sp., Clavibacter* sp.; *Ccal, C. californiensis*; *Csep*, *C. sepedonicus*; *Cneb*, *C. nebraskensis*; *Cins, C. insidiosus*; *Cphas*, *C. phaseoli*; *Czhang, C. zhanzhiyongii*; *Ccap, C. capsici*; *Ctes, C. tessellarius*.

| number | id | Isolation place | Isolation year | Host | Associated paper | DOI |
|---|---|---|---|---|---|---|
| **1** | Ccap RA1B | Mexico | 2017 | tomato (*Solanum lycopersicum*) | this paper | - |
| **2** | Cins CFBP 2404 | USA | 1955 | alfalfa (*Medicago sativa*) | - | - |
| **3** | Cneb 61-1 | USA | 2006 | maize (*Zea mays*) | - | - |
| **4** | Cneb 7580 | USA | 2006 | maize (*Zea mays*) | - | - |
| **5** | Cm CFBP7158 | New Zealand | 1968 | tomato (*Solanum lycopersicum*) | Thapa, 2020 | 10.1094/PHYTO-10-19-0405-R |
| **6** | Cm CFBP5842 | Brazil | 1993 | pepper (*Capsicum annum*) | Thapa, 2020 | 10.1094/PHYTO-10-19-0405-R |
| **7** | Cm CFBP7568 | USA | 2000 | tomato (*Solanum lycopersicum*) | Thapa, 2020 | 10.1094/PHYTO-10-19-0405-R |
| **8** | Cm CFBP7314 | USA | 2002 | tomato (*Solanum lycopersicum*) | Thapa, 2020 | 10.1094/PHYTO-10-19-0405-R |
| **9** | Cm CFBP7315 | USA | 1998 | tomato (*Solanum lycopersicum*) | Thapa, 2020 | 10.1094/PHYTO-10-19-0405-R |
| **10** | Cm ATCC 14456 | Italy | 1961 | tomato (*Solanum lycopersicum*) | Thapa, 2020 | 10.1094/PHYTO-10-19-0405-R |
| **11** | Cm NZ2541 | UK | 1962 | tomato (*Solanum lycopersicum*) | Thapa, 2020 | 10.1094/PHYTO-10-19-0405-R |
| **12** | Cm NT20-18 | the Netherlands | 2020 | tomato (*Solanum lycopersicum*) | this paper | - |
| **13** | Ccap 1207 | South Korea | 1997 | pepper (*Capsicum annum*) | - | - |
| **14** | Cm OP3 | Chile | 2015 | tomato (*Solanum lycopersicum*) | Valenzuela, 2021 | 10.3390/microorganisms9071530 |
| **15** | Cm VL527 | Chile | 2012 | tomato (*Solanum lycopersicum*) | Valenzuela, 2021 | 10.3390/microorganisms9071530 |
| **16** | Cm MSF322 | Chile | 2005 | tomato (*Solanum lycopersicum*) | Valenzuela, 2021 | 10.3390/microorganisms9071530 |
| **17** | Czhang DM1 | Australia | 2017 | barley (*Hordeum vulgare*) | Tian, 2021 | 10.1099/ijsem.0.004786 |
| **18** | C sp LMG 26808 | the Netherlands | unknown | tomato (*Solanum lycopersicum*) | Zaluga, 2014 | 10.1186/1471-2164-15-392 |
| **19** | Cm MX14-E106-3 | Mexico | 2014 | tomato (*Solanum lycopersicum*) | this paper | - |
| **20** | Cm MX13-E96-2 | Mexico | 2013 | tomato (*Solanum lycopersicum*) | this paper | - |
| **21** | Cm MX16-H8B | Mexico | 2016 | tomato (*Solanum lycopersicum*) | this paper | - |
| **22** | Cm MX16-I12A | Mexico | 2016 | tomato (*Solanum lycopersicum*) | this paper | - |

| number | id | Isolation place | Isolation year | Host | Associated paper | DOI |
|---|---|---|---|---|---|---|
| **23** | Cm MX15-L3A | Mexico | 2015 | tomato (*Solanum lycopersicum*) | this paper | - |
| **24** | Cm MX15-115 | Mexico | 2015 | tomato (*Solanum lycopersicum*) | this paper | - |
| **25** | Cm CASJ006 | USA | 2002 | tomato (*Solanum lycopersicum*) | Thapa, 2017 | 10.1094/MPMI-06-17-0146-R |
| **26** | Cm CASJ009 | USA | 2011 | tomato (*Solanum lycopersicum*) | Thapa, 2017 | 10.1094/MPMI-06-17-0146-R |
| **27** | Cm CFBP8019 | the Netherlands | unknown | tomato (*Solanum lycopersicum*) | Thapa, 2017 | 10.1094/MPMI-06-17-0146-R |
| **28** | Cm CFBP7494 | Chile | 1999 | tomato (*Solanum lycopersicum*) | Thapa, 2017 | 10.1094/MPMI-06-17-0146-R |
| **29** | Cm CFBP8017 | the Netherlands | 2006 | tomato (*Solanum lycopersicum*) | Thapa, 2017 | 10.1094/MPMI-06-17-0146-R |
| **30** | Cm CASJ008 | USA | 2002 | tomato (*Solanum lycopersicum*) | Thapa, 2017 | 10.1094/MPMI-06-17-0146-R |
| **31** | Cm Z002 | USA | 2012 | orchard grass (*Dactylis glomerata*) | Davis, 2018 | 10.1128/mBio.01280-18 |
| **32** | Cm AY1B3 | USA | 2014 | perennial ryegrass (*Lolium perenne*) | Davis, 2018 | 10.1128/mBio.01280-18 |
| **33** | Cm AY1A6 | USA | 2014 | perennial ryegrass (*Lolium perenne*) | Davis, 2018 | 10.1128/mBio.01280-18 |
| **34** | Cm AY1B2 | USA | 2013 | perennial ryegrass (*Lolium perenne*) | Davis, 2018 | 10.1128/mBio.01280-18 |
| **35** | Cm Z001 | USA | 2012 | orchard grass (*Dactylis glomerata*) | Davis, 2018 | 10.1128/mBio.01280-18 |
| **36** | Cins R1-1 | USA | 2009 | barrel medic (*Medicago truncatula*) | Lu, 2015 | 10.1128/genomeA.00396-15 |
| **37** | Csep ATCC33113 | Canada | 1968 | potato (*Solanum tuberosum*) | EFSA, 2019 | 10.2903/j.efsa.2019.5670 |
| **38** | Ccap PF008 | South Korea | 1991 | pepper (*Capsicum annum*) | Oh, 2016 | 10.1099/ijsem.0.001311 |
| **39** | Cneb NCPPB 2581 | USA | 1971 | maize (*Zea mays*) | - | - |
| **40** | Cneb DOAB 395 | Canada | 2014 | maize (*Zea mays*) | - | - |
| **41** | Ctes DOAB 609 | USA | 1976 | wheat (*Triticum aestivum*) | - | - |
| **42** | Ctes ATCC 33566 | USA | 1976 | wheat (*Triticum aestivum*) | Li, 2017 | 10.1128/genomeA.00721-17 |
| **43** | Cins LMG 3663 | USA | 1955 | alfalfa (*Medicago sativa*) | Li, 2017 | 10.1128/genomeA.00721-17 |
| **44** | Cins R1-3 | USA | 2009 | barrel medic (*Medicago truncatula*) | - | - |
| **45** | Csep CFIA-Cs3N | Canada | 1976 | potato (*Solanum tuberosum*) | Li, 2017; EFSA, 2019 | 10.1128/genomeA.00721-17; 10.2903/j.efsa.2019.5670 |
| **46** | Csep CFIA-CsR14 | Canada | 1991 | potato (*Solanum tuberosum*) | Li, 2017; EFSA, 2019 | 10.1128/genomeA.00721-17; 10.2903/j.efsa.2019.5670 |
| **47** | Cm NCPPB 382 | UK | 1956 | tomato (*Solanum lycopersicum*) | Gartemann, 2008 | 10.1128/JB.01595-07 |
| **48** | Cm MAI1009 | Uruguay | 2012 | tomato (*Solanum lycopersicum*) | this paper | - |

| number | id | Isolation place | Isolation year | Host | Associated paper | DOI |
|---|---|---|---|---|---|---|
| **49** | Cm 1217 | Russia | 2006 | potato (*Solanum tuberosum*) | - | - |
| **50** | Cm VKM Ac-1790 | Russia | 1993 | Agrostis sp. | Tarlachkov, 2021 | 10.1128/MRA.01400-20 |
| **51** | Cphas VKM Ac-2886 | Russia | 2017 | red elderberry (*Sambucus racemosa*) | Tarlachkov, 2021 | 10.1128/MRA.01400-20 |
| **52** | C sp VKM Ac-2542 | Russia | 1993 | quackgrass (*Elymus repens*) | Tarlachkov, 2021 | 10.1128/MRA.01400-20 |
| **53** | C sp VKM Ac-2872 | USA | 2020 | annual bluegrass (*Poa annua*) | Tarlachkov, 2021 | 10.1128/MRA.01400-20 |
| **54** | C sp VKM Ac-2873 | USA | 2020 | common bent (*Agrostis capillaris*) | Tarlachkov, 2021 | 10.1128/MRA.01400-20 |
| **55** | C sp PvP097 | USA | unknown | switchgrass (*Panicum virgatum*) | - | - |
| **56** | C sp PvP098 | USA | unknown | switchgrass (*Panicum virgatum*) | - | - |
| **57** | C sp PvP036 | USA | unknown | switchgrass (*Panicum virgatum*) | - | - |
| **58** | C sp CFBP 3399 | the Netherlands | 1987 | *Tulipa* sp. | - | - |
| **59** | Cm VQ143 | Chile | 2000 | tomato (*Solanum lycopersicum*) | - | - |
| **60** | Cm VQ28 | Chile | 1996 | tomato (*Solanum lycopersicum*) | - | - |
| **61** | Cm 0317 | USA | 2003 | tomato (*Solanum lycopersicum*) | - | - |
| **62** | Cm CMM84 | Mexico | 2018 | tomato (*Solanum lycopersicum*) | - | - |
| **63** | C sp LMG7333 | Hungary | 1957 | tomato (*Solanum lycopersicum*) | - | - |
| **64** | Cchil CFBP 8217 | Chile | 2007 | tomato (*Solanum lycopersicum*) | - | - |
| **65** | Cphas LPPA 982 | Spain | 2009 | bean (*Phaseolus vulgaris*) | - | - |
| **66** | C sp A6099 | India | 2013 | tomato (*Solanum lycopersicum*) | - | - |
| **67** | Ccal CFBP 8216 | USA | 2000 | tomato (*Solanum lycopersicum*) | - | - |
| **68** | Cm MAI1001 | Uruguay | 2012 | tomato (*Solanum lycopersicum*) | this paper | - |
| **69** | Cm MAI1050 | Uruguay | 2014 | tomato (*Solanum lycopersicum*) | this paper | - |

**Supplementary table 4**: Gene families identified by the pangenomic and enrichment analysis

| Gene family | Present in | Genomic location | Loci name/number | Pfam accesión number | Function (Pfam) | Function (RASTtk) |
|---|---|---|---|---|---|---|
| **1** | Cm subclade | chromosome | PAI | - | - | - |
| **2** | Bcm clade | chromosome | PAI | - | - | - |
| **3** | Cm subclade | chromosome | PAI | - | - | ORF19 |
| **4** | Cm subclade | chromosome | PAI | - | - | - |
| **5** | Cm subclade | chromosome | PAI | - | - | - |
| **6** | Cm subclade | chromosome | PAI | - | - | ParB domain-containing protein nuclease |
| **7** | Bcm clade | chromosome | PAI | PF08843.14 | Nucleotidyl transferase AbiEii toxin, Type IV TA system | - |
| **8** | Bcm clade | chromosome | PAI | PF13338.9 | Transcriptional regulator, AbiEi antitoxin | - |
| **9** | Cm subclade | chromosome | PAI | - | - | putative secreted protein |
| **10** | Cm subclade | chromosome | PAI | - | - | - |
| **11** | Cm subclade | chromosome | PAI | - | - | - |
| **12** | Cm subclade | chromosome | PAI | PF00106.28 | short chain dehydrogenase | Short-chain dehydrogenase |
| **13** | Cm subclade | chromosome | PAI | PF00933.24 | Glycosyl hydrolase family 3 N terminal domain | beta-glucosidase (EC 3.2.1.21) |
| **14** | Cm subclade | chromosome | PAI | PF00528.25 | Binding-protein-dependent transport system inner membrane component | Multiple sugar transport system permease protein |
| **15** | Cm subclade | chromosome | PAI | PF13416.9 | Bacterial extracellular solute-binding protein | ABC transporter, substrate-binding protein (cluster 1, maltose/g3p/polyamine/iron) |
| **16** | Cm subclade | chromosome | PAI | PF00440.26 | Bacterial regulatory proteins, tetR family | Transcriptional regulator, AcrR family |
| **17** | Cm subclade | chromosome | PAI | PF00232.21 | Glycosyl hydrolase family 1 | beta-glucosidase (EC 3.2.1.21) |
| **18** | Cm subclade | chromosome | PAI | PF13377.9 | Periplasmic binding protein-like domain | Transcriptional regulator, LacI family |
| **19** | Cm subclade | chromosome | PAI | PF00331.23 | Glycosyl hydrolase family 10 | Endo-1,4-beta-xylanase (EC 3.2.1.8) |
| **20** | Cm subclade | chromosome | PAI | PF00072.27 | Response regulator receiver domain | Two-component transcriptional response regulator, LuxR family |
| **21** | Cm subclade | chromosome | PAI | PF07730.16 | Histidine kinase | Two-component system sensor histidine kinase |
| **22** | Cm subclade | chromosome | PAI | PF03176.18 | MMPL family | Integral membrane protein |
| **23** | Cm subclade | chromosome | PAI | PF00067.25 | Cytochrome P450 | Putative cytochrome P450 hydroxylase |
| **24** | Cm subclade | chromosome | PAI | PF13370.9 | 4Fe-4S single cluster domain of Ferredoxin I | Ferredoxin-like protein SCO7676 |

| Gene family | Present in | Genomic location | Loci name/number | Pfam accesión number | Function (Pfam) | Function (RASTtk) |
|---|---|---|---|---|---|---|
| **25** | Cm subclade | chromosome | PAI | PF07992.17 | Pyridine nucleotide-disulphide oxidoreductase | Ferredoxin reductase |
| **26** | Cm subclade | chromosome | PAI | PF18120.4 | Domain of unknown function (DUF5597) | beta-galactosidase (EC 3.2.1.23) |
| **27** | Cm subclade | chromosome | PAI | PF04616.17 | Glycosyl hydrolases family 43 | Xylan 1,4-beta-xylosidase (EC 3.2.1.37) |
| **28** | Cm subclade | chromosome | PAI | PF07690.19 | Major Facilitator Superfamily | Uncharacterized MFS-type transporter |
| **29** | Cm subclade | chromosome | PAI | PF17389.5 | Bacterial alpha-L-rhamnosidase 6 hairpin glycosidase domain | alpha-L-rhamnosidase (EC 3.2.1.40) |
| **30** | Outside Bcm | chromosome | - | PF01613.21 | Flavin reductase like domain | - |
| **31** | Bcm clade | chromosome | - | - | - | - |
| **32** | Bcm clade | chromosome | - | - | - | - |
| **33** | Outside Bcm | chromosome | 1 | PF07690.19 | Major Facilitator Superfamily | Uncharacterized MFS-type transporter |
| **34** | Cm subclade | chromosome | 1 | - | - | - |
| **35** | Outside Bcm | chromosome | 1 | PF13411.9 | MerR HTH family regulatory protein | Regulatory protein MerR |
| **36** | Outside Bcm | chromosome | 2 | PF13377.9 | Periplasmic binding protein-like domain | Transcriptional regulator, LacI family |
| **37** | Outside Bcm | chromosome | 2 | PF13416.9 | Bacterial extracellular solute-binding protein | ABC transporter, substrate-binding protein (cluster 1, maltose/g3p/polyamine/iron) |
| **38** | Outside Bcm | chromosome | 2 | PF00528.25 | Binding-protein-dependent transport system inner membrane component | Possible alpha-xyloside ABC transporter, permease component |
| **39** | Outside Bcm | chromosome | 2 | PF01055.29 | Glycosyl hydrolases family 31 | alpha-xylosidase (EC 3.2.1.177) |
| **40** | Outside Bcm | chromosome | - | PF13620.9 | Carboxypeptidase regulatory-like domain | Probable hemagglutinin/hemolysin-related protein |
| **41** | Outside Bcm | chromosome | - | - | - | - |
| **42** | Outside Bcm | chromosome | - | - | - | - |
| **43** | Outside Bcm | chromosome | - | - | - | - |
| **44** | Cm subclade | chromosome | - | - | - | - |
| **45** | Cm subclade | chromosome | 3 | PF11706.11 | CGNR zinc finger | - |
| **46** | Cm subclade | chromosome | 3 | PF02627.23 | Carboxymuconolactone decarboxylase family | 4-carboxymuconolactone decarboxylase (EC 4.1.1.44) |
| **47** | Cm subclade | chromosome | 3 | PF02126.21 | Phosphotriesterase family | - |
| **48** | Cm subclade | chromosome | - | - | - | - |
| **49** | Outside Bcm | chromosome | - | PF00107.29 | Zinc-binding dehydrogenase | Putative oxidoreductase |
| **50** | Cm subclade | chromosome | - | - | - | - |

| Gene family | Present in | Genomic location | Loci name/number | Pfam accesión number | Function (Pfam) | Function (RASTtk) |
|---|---|---|---|---|---|---|
| **51** | Cm subclade | chromosome | - | - | - | - |
| **52** | Cm subclade | chromosome | 4 | PF01636.26 | Phosphotransferase enzyme family | - |
| **53** | Cm subclade | chromosome | 4 | PF13302.10 | Acetyltransferase (GNAT) domain | - |
| **52** | Cm subclade | chromosome | 4 | PF00501.31 | AMP-binding enzyme | Long-chain-fatty-acid--CoA ligase (EC 6.2.1.3) |
| **54** | Cm subclade | chromosome | 4 | PF00109.29 | Beta-ketoacyl synthase, N-terminal domain | 3-oxoacyl-[acyl-carrier-protein] synthase, KASII (EC 2.3.1.179) |
| **55** | Cm subclade | chromosome | 4 | PF02776.21 | Thiamine pyrophosphate enzyme, N-terminal TPP binding domain | - |
| **56** | Cm subclade | chromosome | 4 | PF02441.22 | Flavoprotein | Phosphopantothenoylcysteine decarboxylase (EC 4.1.1.36) homolog |
| **56** | Cm subclade | chromosome | 4 | PF00005.30 | ABC transporter | ABC-type antimicrobial peptide transport system, ATPase component |
| **57** | Cm subclade | chromosome | 4 | PF02687.24 | FtsX-like permease family | ABC transporter, permease protein |
| **58** | Cm subclade | chromosome | 4 | - | - | - |
| **59** | Cm subclade | chromosome | 4 | PF11139.11 | Sap, sulfolipid-1-addressing protein | - |
| **60** | Bcm clade | chromosome | - | - | - | - |
| **61** | Bcm clade | chromosome | - | - | - | - |
| **62** | Outside Bcm | chromosome | - | - | - | - |
| **63** | Bcm clade | chromosome | - | - | - | - |
| **64** | Cm subclade | chromosome | - | - | - | - |
| **65** | Bcm clade | chromosome | - | - | - | - |
| **66** | Outside Bcm | chromosome | - | - | - | - |
| **67** | Outside Bcm | chromosome | - | - | - | - |
| **68** | Bcm clade | chromosome | - | - | - | - |
| **69** | Outside Bcm | chromosome | - | PF08378.14 | Nuclease-related domain | - |
| **70** | Cm subclade | chromosome | - | - | - | - |
| **71** | Cm subclade | chromosome | - | - | - | - |
| **72** | Bcm clade | chromosome | - | - | - | - |
| **73** | Cm subclade | chromosome | - | - | - | - |
| **74** | Cm subclade | chromosome | - | - | - | - |

| Gene family | Present in | Genomic location | Loci name/number | Pfam accesión number | Function (Pfam) | Function (RASTtk) |
|---|---|---|---|---|---|---|
| **75** | Outside Bcm | chromosome | 5 | PF12730.10 | ABC-2 family transporter protein | - |
| **76** | Outside Bcm | chromosome | 5 | PF12730.10 | ABC-2 family transporter protein | putative transporter, trans-membrane domain bacteriocin immunity protein |
| **77** | Outside Bcm | chromosome | 5 | PF00072.27 | Response regulator receiver domain | Two-component transcriptional response regulator, LuxR family |
| **78** | Outside Bcm | chromosome | 5 | PF07730.16 | Histidine kinase | two-component sensor |
| **79** | Outside Bcm | chromosome | 5 | - | - | - |
| **80** | Outside Bcm | chromosome | 5 | PF13575.9 | Domain of unknown function (DUF4135) | Lanthionine biosynthesis protein LanM |
| **81** | Bcm clade | chromosome | - | - | - | Uncharacterized 29.3 kDa protein (ORF92) |
| **82** | Outside Bcm | chromosome | - | - | - | - |
| **83** | Outside Bcm | chromosome | 6 | - | - | - |
| **84** | Cm subclade | chromosome | 6 | - | - | - |
| **85** | Bcm clade | chromosome | 6 | - | - | - |
| **86** | Cm subclade | chromosome | - | PF01823.22 | MAC/Perforin domain | - |
| **87** | Outside Bcm | chromosome | - | - | - | - |
| **88** | Outside Bcm | chromosome | - | PF13310.9 | Virulence protein RhuM family | Putative DNA-binding protein in cluster with Type I restriction-modification system |
| **89** | Outside Bcm | chromosome | - | - | - | - |
| **90** | Outside Bcm | chromosome | 7 | PF00144.27 | Beta-lactamase | Putative esterase |
| **91** | Outside Bcm | chromosome | 7 | PF00251.23 | Glycosyl hydrolases family 32 N-terminal domain | Sucrose-6-phosphate hydrolase (EC 3.2.1.26) |
| **92** | Outside Bcm | chromosome | 7 | PF00528.25 | Binding-protein-dependent transport system inner membrane component | ABC transporter, permease protein 2 (cluster 1, maltose/g3p/polyamine/iron) |
| **93** | Outside Bcm | chromosome | 7 | PF00528.25 | Binding-protein-dependent transport system inner membrane component | ABC transporter, permease protein 1 (cluster 1, maltose/g3p/polyamine/iron) |
| **94** | Outside Bcm | chromosome | 7 | PF01547.28 | Bacterial extracellular solute-binding protein | ABC transporter, substrate-binding protein (cluster 1, maltose/g3p/polyamine/iron) |
| **95** | Outside Bcm | chromosome | 7 | PF13377.9 | Periplasmic binding protein-like domain | Transcriptional regulator, LacI family |
| **96** | Outside Bcm | chromosome | - | - | - | - |
| **97** | Outside Bcm | chromosome | - | PF01425.24 | Amidase | Allophanate hydrolase (EC 3.5.1.54) |
| **98** | Bcm clade | chromosome | - | - | - | - |
| **99** | Cm subclade | chromosome | 8 | - | - | - |
| **100** | Cm subclade | chromosome | 8 | PF00005.30 | ABC transporter | ABC transporter, ATP-binding protein |

| Gene family | Present in | Genomic location | Loci name/number | Pfam accesión number | Function (Pfam) | Function (RASTtk) |
|---|---|---|---|---|---|---|
| **101** | Cm subclade | chromosome | 8 | - | - | - |
| **102** | Cm subclade | chromosome | 8 | - | - | - |
| **103** | Bcm clade | chromosome | - | - | - | - |
| **1** | Cm subclade | pCM2 | - | - | - | - |
| **7** | Bcm clade | pCM2 | - | PF08843.14 | Nucleotidyl transferase AbiEii toxin, Type IV TA system | - |
| **8** | Bcm clade | pCM2 | - | PF13338.9 | Transcriptional regulator, AbiEi antitoxin | - |
